# From Belief to Brain: How Growth Mindset Optimizes Cortico-Striatal Dynamics for Cognitive Development

**DOI:** 10.1101/2022.07.11.499525

**Authors:** Yuyao Zhao, Junjie Cui, Ze Zhang, Hui Zhao, Jiahua Xu, Menglu Chen, Lei Hao, Ying He, Hui Wang, Yanpei Wang, Daoyang Wang, Yueqing Hu, Zhuo Rachel Han, Shuping Tan, Weiwei Men, Jiahong Gao, Lang Chen, Yong He, Sha Tao, Qi Dong, Shaozheng Qin

## Abstract

Growth mindset—the belief that abilities are malleable through effort—drives motivation, action and achievement. Yet, the underlying mechanisms remain elusive, necessitating a unified framework that integrates cognitive, neural, and developmental processes. Leveraging longitudinal neuroimaging and computational modeling to reveal moment-to-moment decision responses and brain state dynamics during working memory (WM), we show that growth mindset enhances WM development from middle childhood to adolescence via nuanced cortico-striatal network dynamics that balance between brain state flexibility and stability. Behaviorally, children with higher growth mindset exhibited better WM performance both cross-sectionally and longitudinally, attributed to faster evidence accumulation during moment-to-moment information updating, especially under high task demands. Neurally, a higher growth mindset was associated with greater activation of the dorsal striatum, cingulo-opercular (action-mode), and fronto-parietal networks during WM, which further accounted for longitudinal WM improvement and latent evidence accumulation. Such cortico-striatal activation covaried with neurochemical mediators critical for motivation and executive functioning. Analyzing non-stationary network dynamics revealed that growth mindset optimized dynamical organization of cortico-striatal networks, with an activated state highly flexible to support moment-to-moment information updating and a deactivated state remaining stable across blocks to suppress irrelevant information. This suggests a balanced allocation of resources for accumulating evidence while suppressing noise during WM. Together, our findings support a neurocognitive framework in which growth mindset enhances WM development via nuanced orchestration of cortico-striatal networks to enable efficient dynamic computations and foster far-reaching cognitive development.

## Introduction

Growth mindset refers to the underlying beliefs that people have about their abilities and skills being malleable and improvable through continuous efforts. Individuals who endorse a growth mindset often demonstrate higher performance in a spectrum of cognitive tasks, skills, academic and professional achievements(*1–4*). Notably, growth mindset interventions are recognized to enlighten latent potentials in school-aged children(*2*, *5–8*), whose rapidly maturing brains and evolving cognitive abilities are plastic and receptive to educational influences. Targeting this critical developmental window, growth mindset has been shown to yield robust benefits in children’s math learning and executive functions(*1*, *6*, *9*, *10*). Such benefits are theorized to foster motivation and executive functions, which optimize regulation of neurocognitive resources in achieving goal-directed behaviors(*11–13*). Despite decades of research and application in educational practice, the underlying neurocognitive mechanisms remain fragmented. Even less is known about how such benefits evolve over development in younths. A unified model integrating cognitive, neural, and developmental processes is crucial to advance our understanding of the underlying mechanisms of growth mindset’s merits.

Several theoretical models have attempted to account for the merits of growth mindset. Dweck’s mindset theory contrasts growth mindset with fixed mindset - the belief that abilities are static and unmalleable through effort. Challenges are appraised as opportunities, engaging reward systems rather than threats. Students with growth mindset persevere through academic challenges, and professionals embrace feedback and adapt accordingly(*14*, *15*). Neuroscientific evidence also supports the brain’s capacity for change, reinforcing the theory’s core premise(*6*, *16*, *17*). The self-determination theory posits that the perception of self-control and mastery can initiate and maintain motivation, driving individuals to pursue goal-directed behaviors and achieve higher performance(*18*, *19*). Likewise, adopting a growth mindset fosters intrinsic motivation, such as the desire to learn or improve for personal satisfaction, thereby enhancing engagement in cognitive tasks requiring sustained effort toward long-term goals(*20–22*). Recent neurocognitive models link growth mindset to adaptability in cognitive strategies and executive functions anchored onto large-scale brain networks(*23*, *24*). Individuals with growth mindset embrace more flexible attention allocation for goal-directed information, while suppressing irrelevant distractions(*3*, *11*, *13*, *25*), and they tend to accumulate and integrate environmental feedback more effectively, utilizing this information for decision-making and task execution(*3*, *11*). Although widely studied, longitudinal evidence remains scarce regarding how growth mindset fosters developmental improvement in executive functioning among elementary school-aged children. Crucially, the neurobiological mechanisms through which growth mindset modulates the developing brain’s cognitive and motivational systems—thereby supporting cognitive development—are poorly understood. Resolving these gaps necessitates a dual approach: integrating cross-sectional neural correlates with longitudinal behavioral trajectories across critical phases of elementary school.

Recent advances in computational modeling of moment-to-moment decision responses enable us to identify latent dynamic computations in various cognitive domains(*26*, *27*). The N-back task, analogous to speeded decision-making, can be modeled as an evidence accumulation process during which effective information extracted from a stream of noisy inputs is rapidly accumulated until sufficient evidence reaches the threshold to make a decision, at which point a response is executed(*28–30*). The Hierarchical Drift Diffusion Model (HDDM) has been widely used to decompose decision responses in a given task into latent decision-making dynamics modulated by free parameters. Of these parameters, the speed of evidence accumulation refers to “drift rate”, reflecting the ability to extract effective information from perceived inputs(*26*). As such, the HDDM allows us to decipher which latent processes involving deliberate dynamic decision-making would be most augmented by growth mindset. Thus, it is conceivable that growth mindset would enhance cognitive performance in school-aged children, most likely via acting on latent evidence accumulation during decision-making dynamics.

The mastery of growth mindset involves multiple brain systems and networks. Growth mindset-enhanced cognitive performance has been linked to greater activation in core regions of the frontoparietal network (FPN), cingulo-opercular network (CON) (core part of the action-mode network, AMN), and striatal systems critical for executive functions and motivation(*3*, *6*, *11*, *13*). Recent studies also emphasize that cortico-striatal connectivity, particularly between the anterior cingulate cortex (ACC) and striatum, plays a critical role in improving math skills among individuals who endorse growth mindset(*6*, *31*). Notably, the dorsal rather than ventral striatum engages more in selecting, filtering, and updating of input information during working memory(*32*, *33*), which can be actively shaped by dopaminergic projections(*34–36*). These findings provide insights into the neural bases of growth mindset, but the neurocognitive pathways of how these systems are coordinated to enable growth mindset-fostered longitudinal improvement in school-aged children remain elusive. Based on neuroimaging observations, we hypothesize that growth mindset would enhance children’s cognitive performance by increasing engagement of core nodes in cortico-striatal networks that may covary with dopaminergic modulators. Such effects at earlier ages would transform into longitudinal improvement over development according to the self-determination theory.

The neurobiology of motivation-cognition interactions has recognized an intricate interplay of the prefrontal cortex and striatal circuitry in support of goal-directed behaviors(*37*, *38*). The dynamic assembly of cortico-striatal regions, with activated and deactivated states, is critical both cognitive and motivational processes to support optimal performance in effortful tasks(*39*, *40*). Motivation-driven dopamine releases, by acting on prefrontal and dorsal striatal circuits(*34–36*), can energize and facilitate flexible allocation of neural resources in response to changing task demands, as well as evaluation and filtering of external information to enable efficient processing(*33*, *41*, *42*). Core nodes of the FPN, such as the dorsolateral prefrontal cortex, are crucial for evidence accumulation during information updating and cognitive control pertinent to goal-directed behavior(*43*, *44*). Functional coupling and decoupling among the FPN, AMN and striatal regions are crucial for initiating and maintaining a high-arousal “action-mode” in response to external task demands(*45*). The ACC is crucial for monitoring and integrating information, thereby enhancing cognitive flexibility and adaptability(*42*, *46*, *47*). Dynamic network modeling such as Hidden Markov modeling (HMM) offers a useful approach to identify time-resolved functional brain network (re)configurations reflecting latent brain states at each time point involved in a given task(*48*, *49*). Based on Viterbi decoded sequence, brain state dynamics can be quantified by fractional occupancy and system-level state transitions(*50*), providing an ideal approach to probe how growth mindset modulates cortico-striatal network dynamics to support cognitive performance. According to above mindset theories and network dynamics, we hypothesize that growth mindset would optimize cortico-striatal network dynamics with flexibility and stability of distinct brain states to enable a balanced allocation of neural resources for accumulating evidence during moment-to-moment information updating, while maintaining a stable goal for each task demand.

To test above hypotheses, we here leverage functional magnetic resonance imaging (fMRI), in conjunction with computational modeling of trial-by-trial decision responses during N-back WM task, to investigate how growth mindset fosters longitudinal WM development in 454 school-aged children (8-12 years old at baseline) over three years (**Figure 1A**). Three WM loads were set to obtain growth mindset effects on task demands and task-invoked brain responses. To probe the longitudinal effects of growth mindset on WM development, children were invited back for follow-up measurements each year for WM assessments. The HDDM was implemented to estimate key parameters reflecting latent computational dynamics during WM processing, including drift rate and decision threshold. Brain-wide activation and multiple regression were employed to identify brain systems linked to growth mindset and WM performance. Moreover, meta-analytic decoding and neurochemical approaches were used to further identify mental processes corresponding to our identified brain systems. Finally, the HMM was used to probe the dynamic organization of large-scale cortico-striatal networks reflecting brain state dynamics and flexible resource allocation. Structural equation modeling was then used to address how cortico-striatal coactivation and network dynamics contribute to cross-sectional and longitudinal effects of growth mindset on developmental improvement in cognitive performance.

**Figure 1.**
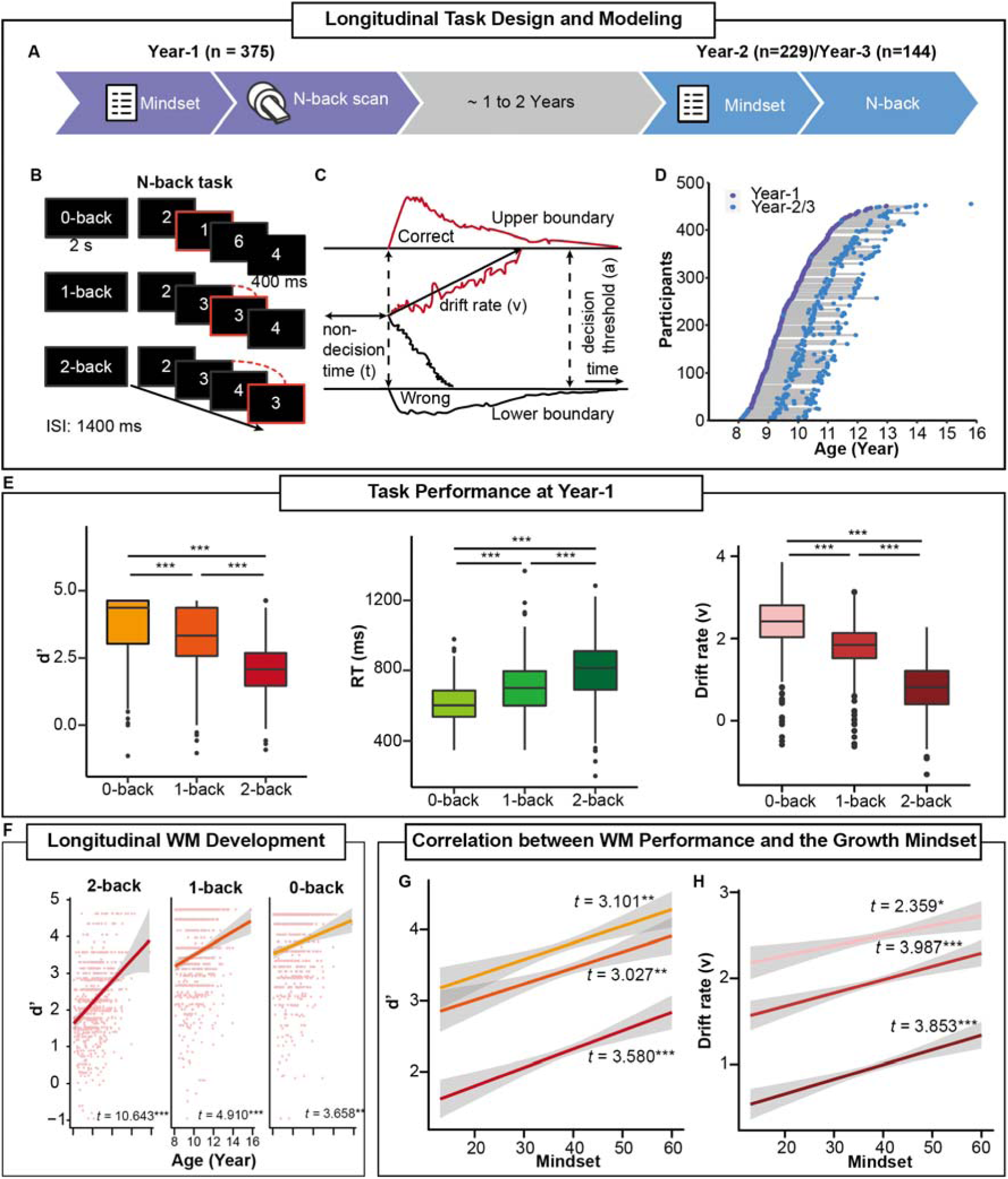
Growth mindset fosters WM development over three years in children. **(A)** Longitudinal design to investigate the effects of growth mindset on WM development over three years and its underlying neurocognitive substrates. Both behavioral and fMRI data were obtained in Year-1. A subsample of children was invited back for growth mindset and N-back tasks in Year-2 and/or Year-3. (**B)** A schematic view of the numerical N-back task with three WM loads. **(C)** An illustration of the HDDM with three free parameters: drift rate (v) indicates the speed of evidence accumulation, decision threshold (a) is the boundary for decision choice, and non-decision time (t) reflects other processes like early stimulus encoding. The red and black lines represent the correct and incorrect decision paths, respectively. **(D)** Age distribution of children at each visit. Each dot represents one child at the time of tests. Dots of the same child are connected with lines. **(E)** Boxplots of discrimination indices (d’), reaction time (RTs), and drift rate (v) during WM in Year-1. The thick black line in each box represents the median, with the 25th and 75th percentiles**. (F)** Developmental trajectories of WM performance measured by d’. Each dot represents one observation. **(G)** Growth mindset positively associated with WM d’ across three years (total observation = 748). Different shades of orange or red represent WM workloads. **(H)** Growth mindset positively correlated with drift rate across three years. Age and gender were controlled for regressions. Shading represents a 95% confidence interval (CI). Notes: ***p < 0.001, **p < 0.01, Bonferroni-corrected.

## Results

### Growth mindset enhances both cross-sectional and longitudinal WM development

We first examined whether growth mindset promotes children’s WM performance throughout development from 8 to 16 years. As presented in **Figure 1E**, children responded more slowly and with less accuracy under high-load conditions, whereas the performance improved with age (**Figure 1F**). More detailed WM-loading manipulations and developmental trajectories of all measures are provided in the **Supplementary Materials**. In the first visit (Year-1), growth mindset was positively correlated with WM performance measured by d’ and reaction times (RTs), with the most prominent effect observed in the 2-back condition (0-back: *r* = 0.194, *p* < 0.001; 1-back: *r* = 0.134, *p* = 0.010; 2-back: *r* = 0.213, *p* < 0.001; all corrected; **Figure S2**). Mixed-effect model analysis of longitudinal data, with age and gender as covariates, revealed that growth mindset predicted children’s longitudinal WM performance (d‘) over three years, especially in the 2-back condition (β = 0.122, *t* = 3.580, *p* < 0.001, corrected), as well as in the 0- and 1-back conditions (both β > 0.119, *t* > 3.027, *p* < 0.003, corrected) (**Figure 1G, Table S9**). However, we did not observe age-related interaction effects between growth mindset and WM performance (β = -0.011, *t* = -0.332, *p* = 0.740), indicating that children’s growth mindset remains stable throughout development.

We then implemented the HDDM to unravel the benefits of growth mindset and to examine how latent cognitive dynamics are involved in WM task. Model comparisons revealed that the three parameters of drift rate (v), decision threshold, and non-decision time constituted the best-fitting model (**Figure 1C & S3, Table S2**). Among these parameters, drift rate emerged to show a positive correlation with growth mindset in the first visit (0-back: *r* = 0.132, *p* =0.010; 1-back: *r* = 0.157, *p* = 0.002; 2-back: *r* = 0.193, *p* < 0.001) and over three years later (0-back: β = 0.059, *t* = 2.359, *p* = 0.019; 1-back: β = 0.091, *t* = 3.987, *p* < 0.001; 2-back: β = 0.087, *t* = 3.853, *p* < 0.001; **Figure 1H, Table S10**), with the most prominent effect in the 2-back condition. These results indicate the benefits of growth mindset on children’s WM development spanning over three years, with the most prominent effect observed under high task demand (2-back) condition and attributed to improvements in latent evidence accumulation during information updating in WM task.

### Growth mindset is associated with greater activity in cortico-striatal systems during WM

Next, we aimed to identify the neural correlates underlying the benefits of growth mindset on children’s WM performance over development. A whole-brain multiple regression analysis was conducted for WM-related neural activity maps derived from the contrast of 2-versus 0-back condition, with children’s growth mindset as a covariate of interest at the first visit (Year-1) while controlling for age and gender. This analysis revealed significant clusters in the dorsal striatum, cingulo-opercular, and fronto-parietal regions (**Figure 2A&B, Table S11**) critical for executive functions based on meta-analytic mapping (**Figure 2C**). More importantly, we observed that children’s growth mindset was positively associated with greater activation in the dorsal striatum during WM, especially located at the dorsolateral caudate **(Figure 2B)**. We also observed positive correlations between children’s growth mindset and WM-related activation in core nodes of the cingulo-opercular network (CON) and frontal-parietal network (FPN). The CON here includes the supplemental motor area (SMA) extending into dorsal anterior cingulate cortex (dACC) and right anterior insula (aIns); the FPN includes the dorsolateral prefrontal cortex (dlPFC), inferior parietal sulcus (IPS), and frontal eye field (FEF). In addition, an opposite correlation was observed in the posterior cingulate cortex (PCC) and parahippocampus (PHC). A similar pattern of results was observed for the contrast map of the 2-versus 1-back condition, but null effects were observed for the contrast map between the 1- and 0-back conditions.

**Figure 2.**
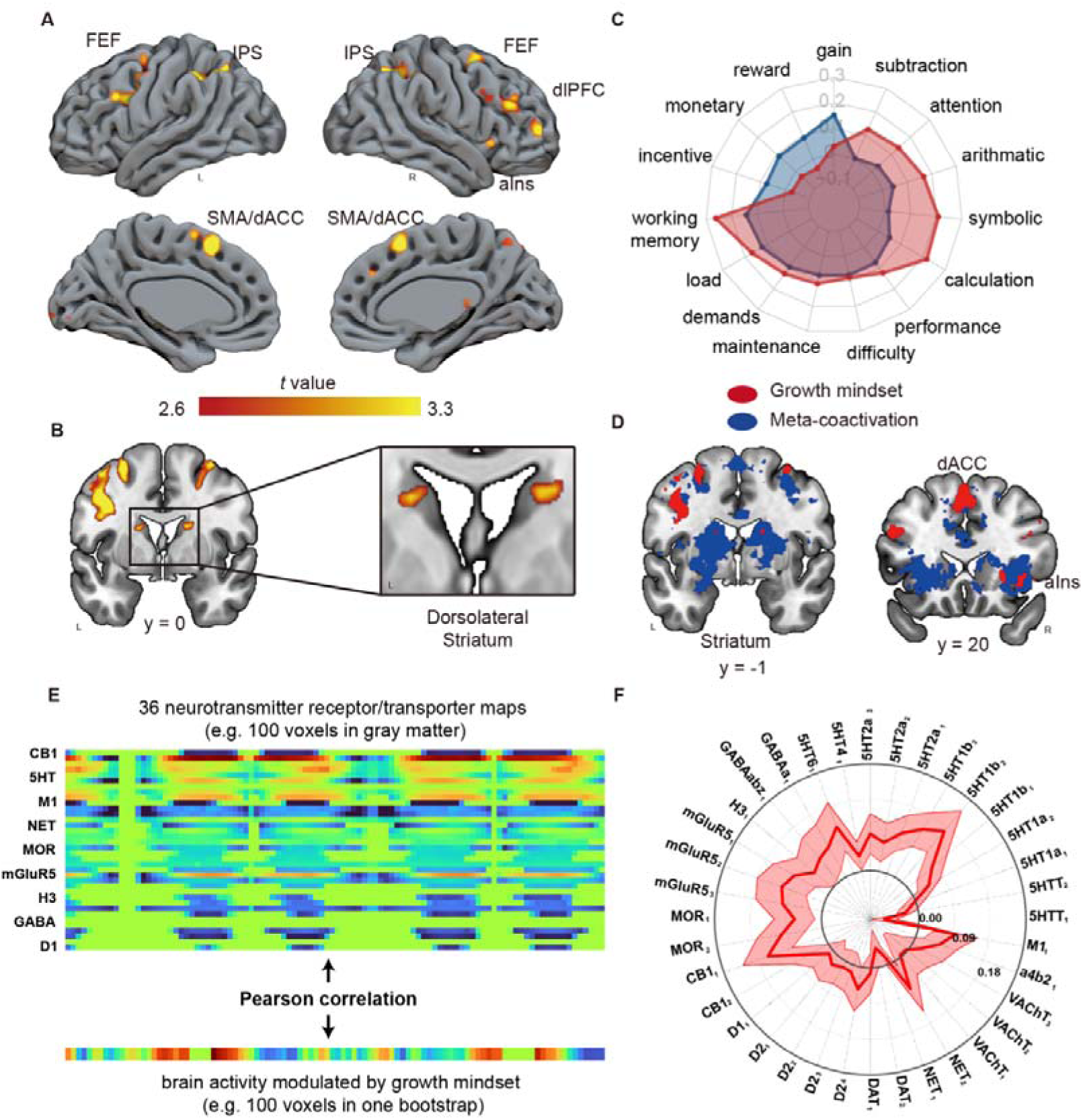
Brain systems associated with children’s growth mindset under high WM demand. **(A, B)** Significant clusters from multiple regression analysis for the contrast map of 2-with 0-back condition show positive correlations with children’s growth mindset at Year-1. (**C**) Meta-analytic decoded terms showing highest correlation with neural response of growth mindset (red) and coactivation (blue) map with the dorsolateral caudate. Correlation (r) in the polar chart depicts the similarity between term-related neural response and decoded map.(**D**) Overlap of children’s growth mindset-related brain regions (red) and meta-analytic coactivation of the dorsolateral caudate (blue). (**E**) An illustration of neurotransmitter analysis. The neurochemical characteristics of growth mindset-related brain map were annotated according to its spatial similarity with 36 independent neurotransmitter receptor/transporter maps. Bootstraps of multiple regression analysis for the contrast map of 2-with 0-back condition were complemented for error estimation. (**F**) Pearson correlations between our identified growth mindset-related brain activity map and neurochemical binding maps. Higher correlation coefficients indicate greater similarity between the two maps. The red shadow illustrates the standard errors of correlation coefficients. The neurotransmitter system can be considered as significant correlations with growth mindset-related brain activity map if the CI is beyond zero. Notes: dlPFC: dorsal lateral prefrontal cortex, FEF: frontal eye field, aIns: anterior insula, IPS: inferior parietal sulcus, dACC: dorsal anterior cingulate cortex. Neurotransmitter systems include cannabinoids (CB1 receptor), dopamine (receptors D1, D2, and transporter), norepinephrine (NET), acetylcholine (VAChT), nicotinic (a4b2, and muscarinic M1), serotonin (5HTT transporter, and 5HT receptors 1a, 1b, 2a, 4, and 6), GABA (GABA-A, GABA-aBZ receptor), histamine (H3), glutamate (metabotropic receptor mGluR5), and opioids (mu-opioid receptor, MOR)).

We then applied a meta-analytic decoding approach to characterize which psychological processes are most likely associated with growth mindset-related brain activity in the cortico-striatal systems from the large-scale Neurosynth platform with over ten thousand neuroimaging studies. Using our identified dorsal striatum as a seed, we first obtained a coactivation map associated with this seed from the Neurosynth platform (**Figure 2C & 2D**) and then compared it with the children’s growth mindset-related activity map from our present study (**Methods**). These two maps were highly overlapped, indicating that our observed cortical activation in SMA/dACC, aIns, IPS, dlPFC, and FEF is indeed often covaried with the dorsal striatum **(Figure 2D; Table S14)**. A meta-analytic decoding approach revealed that children’s growth mindset-related neural activity and dorsal striatal coactivation are associated with terms of executive functions, including WM, calculation, etc., as well as motivation-related processes, including “gain”, “reward”, “monetary” (**Figure 2C**).

To explore whether children’s growth mindset-related brain activity under high WM demand covaries with engagement of motivation-related brain circuitry, we further annotated the neurochemical characteristics linked to the neurotransmitter receptor/transporter maps derived from an open-source PET dataset. A set of spatial similarity metrics was computed for children’s growth mindset-related brain activity covarying with nine different neurotransmitter maps (**Figure 2E**). As shown in **Figure 2F**, though all spatial similarity metrics were significant based on 95% CI of bootstraps, we observed the highest similarity of growth mindset-related activity map with cannabinoids (CB1), glutamate (metabotropic receptor mGluR5), and serotonin (5HT1b2) critical for motivation and reward processing. Together, these results indicate that children’s growth mindset is associated with greater WM-related activation in core regions of the cortico-striatal networks and motivation-related neurotransmitter activity patterns critical for motivational and executive functioning.

### Growth mindset fosters WM development via greater activity in cortico-striatal systems

To investigate the relationships among children’s growth mindset, WM-related brain activity, WM performance, and latent decision-making dynamics, we then restrained our analysis on core regions of interest (ROIs) in the striatum, CON, FPN, and DMN defined by an independent Neurosynth meta-analysis (**Figure 3A & S4**). As we expected, this analysis revealed that task-invoked activity in these ROIs was positively correlated with children’s growth mindset with cross-sectional and longitudinal WM performance and three latent HDDM parameters, while including age and gender as covariates of no interest (**Table S15-16**). Specifically, WM-related activity in the dorsal striatum (*r* = 0.150, p = 0.014, corrected) and CON regions (aIns *r* = 0.167, SMA/dACC: *r* = 0.201; all *p* ≤ 0.007, corrected) was positively correlated with children’s growth mindset and faster drift rate, especially under the 2-back condition (all r ≥ 0.172, all *p* ≤ 0.004, corrected). Task-invoked activity in FPN regions, including dlPFC, FEF, and IPS, was also positively associated with children’s growth mindset (all *r* ≥ 0.181, all *p* ≤ 0.004, corrected) and faster drift rate (all *r* ≥ 0.124, all *p* ≤ 0.034, corrected). Among DMN regions, WM-related activity in these regions was negatively correlated with drift rate (vmPFC *r* = -0.224, PCC *r* = -0.151; HPC/PHC: *r* = -0.157, all *p* ≤ 0.011, corrected) except the AG (*r* = -0.096, *p* = 0.096).

**Figure 3.**
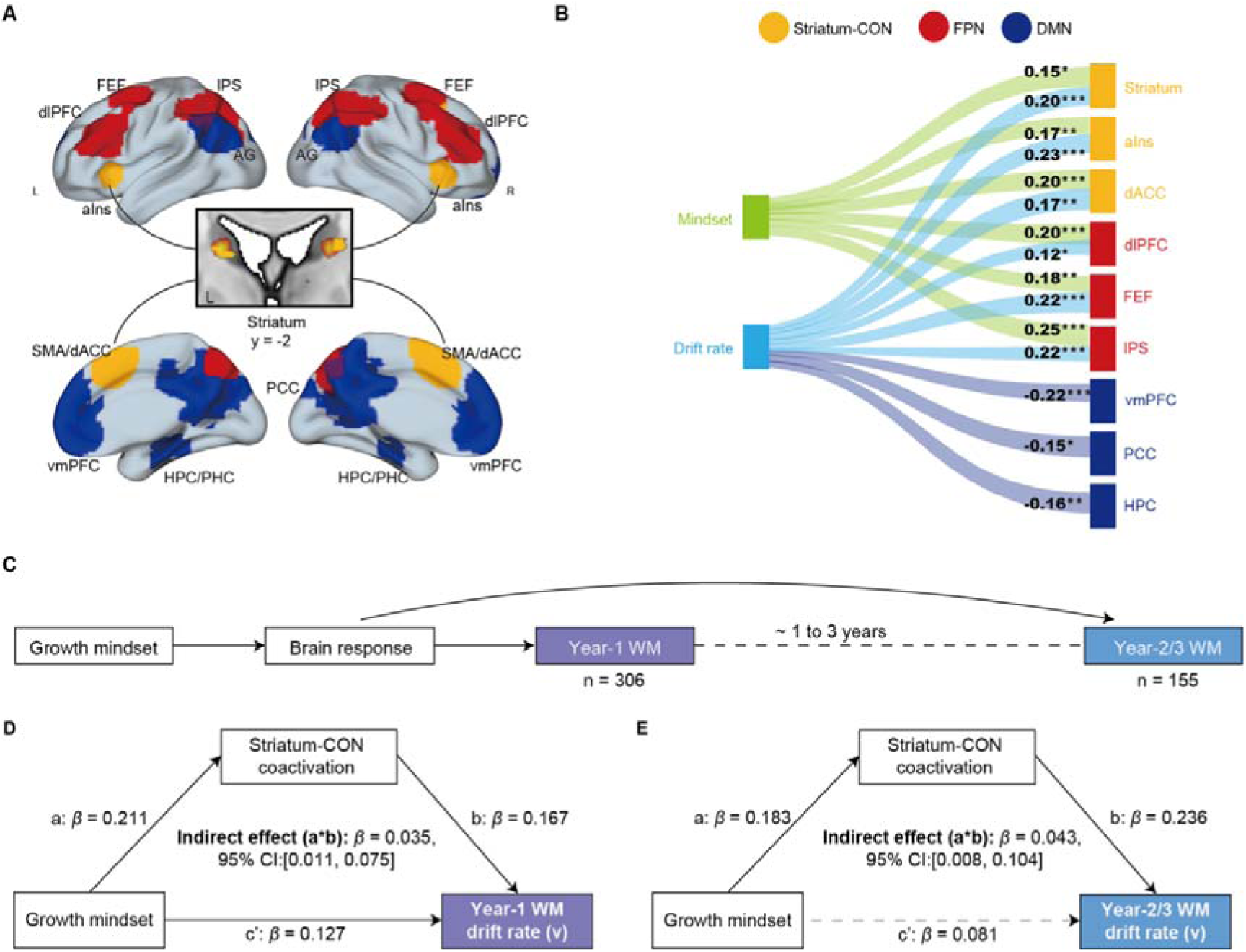
Growth mindset improves both cross-sectional and longitudinal WM drift rates via striatal and CON network coactivation. **(A)** ROIs are defined for the striatum-CON (yellow), FPN (red), and DMN (blue). Task-invoked response under 2- vs. 0-back conditions were exacted from each ROI for further analysis. (**B)** Flow diagram of correlations between activity in each ROI and growth mindset (and drift rate) at first visit. Only the drift rate in the 2-back condition was illustrated here regarding its high correlation with growth mindset compared to the 1-back condition. **(C)** An illustration of cross-sectional and longitudinal data analysis. The mediatory role of growth mindset-related ROIs was examined in both current (Year-1) and future (Year-2/3) performance. **(D, E)** Mediation models of task-invoked activity in striatum-CON regions accounting separately for cross-sectional and longitudinal benefits of growth mindset on WM drift rate. Task-invoked activation in cortico-striatal regions during 2-versus 0-back condition could account for the beneficial effects of children’s growth mindset on higher drift rate at first visit as well as longitudinal outcomes. Notes: Age and gender were set as covariances, and all p values were corrected for multiple comparisons using FDR correction. ***p < 0.001, **p < 0.01, *p < 0.05, n.s.: no significance

To test our hypothesis about how children’s growth mindset improves WM development through cortico-striatal functional organization, we implemented structural equation modeling (SEM) to investigate potential mediation pathways among children’s growth mindset, cross-sectional and longitudinal WM performance, and drift rate via WM-related activity in cortico-striatal regions. We examined both cross-sectional and longitudinal mediatory effects of the striatum-CON while also testing the mediatory role of FPN as indicated by the above findings. It showed that the coactivation of both the striatum-CON system and FPN mediated the relationship between growth mindset and WM performance (d’) at first visit (indirect Est. = 0.039, 95% CI = [0.014, 0.077]; **Figure 3C, D & E; Table S17**). A similar mediatory effect was also observed in the FPN regions (indirect Est. = 0.043, 95% CI = [0.015, 0.085]). The coactivation of striatum-CON and FPN regions can also account for the beneficial effects of growth mindset on drift rate at first visit (striatum-CON: indirect Est. = 0.035, 95% CI = [0.011, 0.075]; FPN: indirect Est. = 0.047, 95% CI = [0.019, 0.089]) (Figure 3D; Table S18).

More importantly, for the question at issue, we further investigated whether and how task-invoked activation in striatum-CON and FPN regions accounts for the benefits of children’s growth mindset on longitudinal WM performance and drift rate (reflecting latent evidence accumulation during information updating) across three years. Although we observed null direct effects of children’s growth mindset on longitudinal WM performance (d’: *r* = 0.015, RTs: *r* = 0.033, both *p* > 0.69), task-invoked activity in SMA/dACC at first visit (*r* = 0.191, *p* = 0.019) was positively associated with growth mindset, which could further account for longitudinal improvement in WM performance with faster RTs (*r* = -0.193, *p* = 0.018) and higher drift rate (*r* = 0.199, *p* = 0.014) across three years, even after controlling for performance at baseline. Such associations only emerged in the SMA/dACC response (**Table S19 & S20**). Though task-invoked responses in FPN regions at first visit were associated with faster RTs in longitudinal WM assessment (FEF: *r* = -0.284, *p* < 0.001; IPS: *r* = -0.257, *p* = 0.002), they were not significantly correlated with children’s growth mindset (FEF: *r* = 0.101, *p* = 0.219; IPS: *r* =0.148, *p* = 0.070). Further mediation analysis of longitudinal data revealed that task-invoked response in the striatum-CON system at the first visit could account for the indirect association between children’s growth mindset and longitudinal improvement in WM performance across three years (indirect Est. = 0.034, 95% CI = [0.005, 0.090]; **Table S21**). Parallel analysis revealed that task-invoked activation in the striatum-CON systems could also account for the indirect association between children’s growth mindset and longitudinal improvement in drift rate (indirect Est. = 0.043, 95% CI = [0.008, 0.104]; **Figure 3E; Table S22**). These results indicate that striatum-CON systems play a mediatory role in supporting longitudinal improvement in WM performance in youths, especially for the speed of latent evidence accumulation.

### Growth mindset fosters WM development via nuanced cortico-striatal network dynamics with distinct tempospatial state flexibility and stability

Beyond task-invoked regional activity in cortico-striatal networks, we further investigated how these large-scale brain networks are dynamically organized to support WM processing and account for the benefits of children’s growth mindset on cross-sectional and longitudinal WM outcomes. BOLD-fMRI time series of 11 ROIs in cortico-striatal networks linked to children’s growth mindset were first extracted, and we then implemented the HMM to identify latent brain state dynamics during WM processing (**Figure 4A & B**). This analysis revealed a set of eight brain states, each with a unique spatial-temporal configuration of cortico-striatal regions. By evaluating the likelihood of each state occurring at a given time, we were able to identify the most dominant state at each time point **(Figure 4C)**. The distribution of eight brain states aligned well with three WM loads manipulated in our study, demonstrating the effectiveness of our model in decoding three task demands **(Figure 4D&E, S5-7)**.

**Figure 4.**
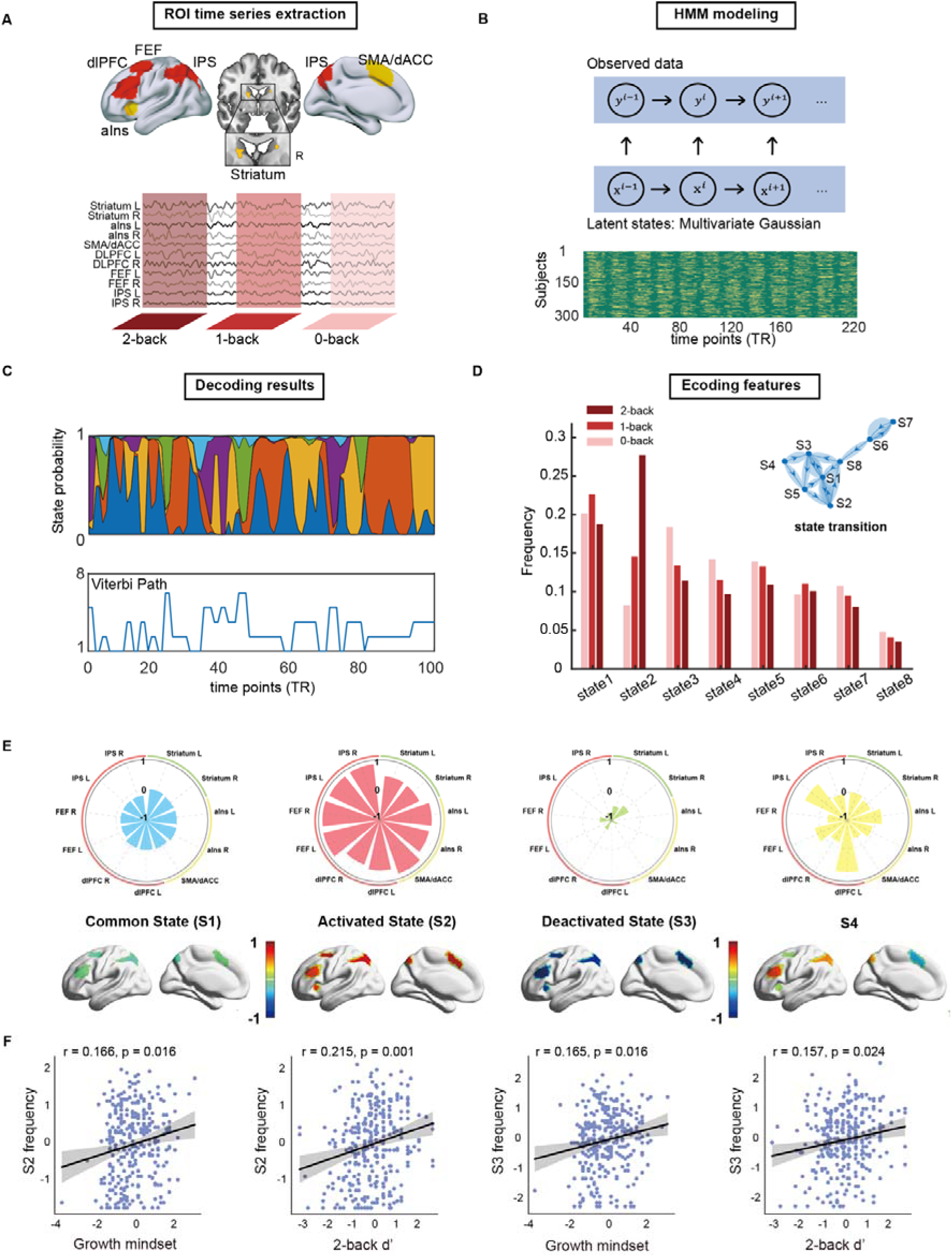
Cortico-striatal network dynamics with the top four latent states under WM tasks. (**A**) The schematic plot of the time series extraction process. Time series from key nodes of the striatum-CON and FPN were extracted to estimate the hidden network dynamics under varied workloads. (**B**) The schematic diagram for HMM estimation. (**C**) The probability of each decoded state as a function of each time point during the entire task. The Viterbi path depicts the most likely sequence of hidden states. (**D**) The frequency of each hidden state under each workload. (**E**) The spatial pattern of the top four states with the highest frequency throughout the task. The value of each area represents the relative activity magnitude, referring to the mean value. (**F**) Regression plots between state frequency and behavioral indexes. Shading represents 95% CI. Each dot represents one subject’s score on the first test.

It is worth noting that State 2 exhibited the highest frequency in high (2-back) than low WM demand (F(2,912) = 167.15, P < 0.001), characterized by greater activation across all cortico-striatal regions. At the same time, State 3 was much more prevalent in low (0-/1-back) than high WM task demand (F(2,912) = 46.57, P < 0.001), with generally lower activation even deactivated across cortico-striatal regions. Thus, States 2 and 3 (as relative to other states) most likely reflect the dynamic organization of cortico-striatal networks, respectively, involved in high and low task demand conditions. Critically, we observed significantly positive correlations of State 2’s and State 3’s frequency with both children’s growth mindset (both r ≥ 0.165, p = 0.016, corrected), as well as WM 2-back performance (both r ≥ 0.157, p ≤ 0.024, corrected) (**Figure 4F**; **Table S23**). These observations indicate that children’s growth mindset is associated with better WM performance and a higher occupancy rate of latent brain states 2 and 3, characterized by generally highly activated and low or even deactivated patterns across regions in the cortico-striatal networks during WM processing.

To understand the temporal dynamics of the above two dominant states, we computed two metrics to quantify temporal flexibility and stability during WM processing (**Figure 5A**; **Methods**). Temporal stability assesses the similarity of a given state’s occurrence between different blocks in each WM 2-back condition. This metric reflects how stable the same state is engaged in different blocks, as it may fluctuate among blocks during the task. Temporal flexibility summarizes the variances of a given state’s probability over time within each block of WM condition, depicting how this state flexibly responds to target and non-target stimuli during WM processing. Correlation analyses further revealed that temporal flexibility of State 2 during WM 2-back was positively correlated with children’s growth mindset (*r* = 0.191, *p* = 0.006), as well as WM performance characterized by d’ (*r* = 0.253, *p* < 0.001) and latent drift rate (*r* = 0.218, *p* = 0.001). Likewise, the temporal stability of State 3 across blocks during the WM 2-back condition was positively correlated with children’s growth mindset (*r* = 0.161, *p* = 0.040), as well as WM d-prime (*r* = 0.306, *p* < 0.001) and drift rate (*r* = 0.283, *p* < 0.001). Notably, these positive correlations are exclusive to States 2 and 3. Further mediation analyses revealed that the beneficial effects of children’s growth mindset on drift rate at the first visit could be separately accounted for by temporal flexibility of State 2 (indirect Est. = 0.037, 95% CI = [0.012, 0.076]; **Figure 5C**) and stability of State3 (indirect Est. = 0.042, 95% CI = [0.016, 0.086]; **Figure 5D**).

**Figure 5.**
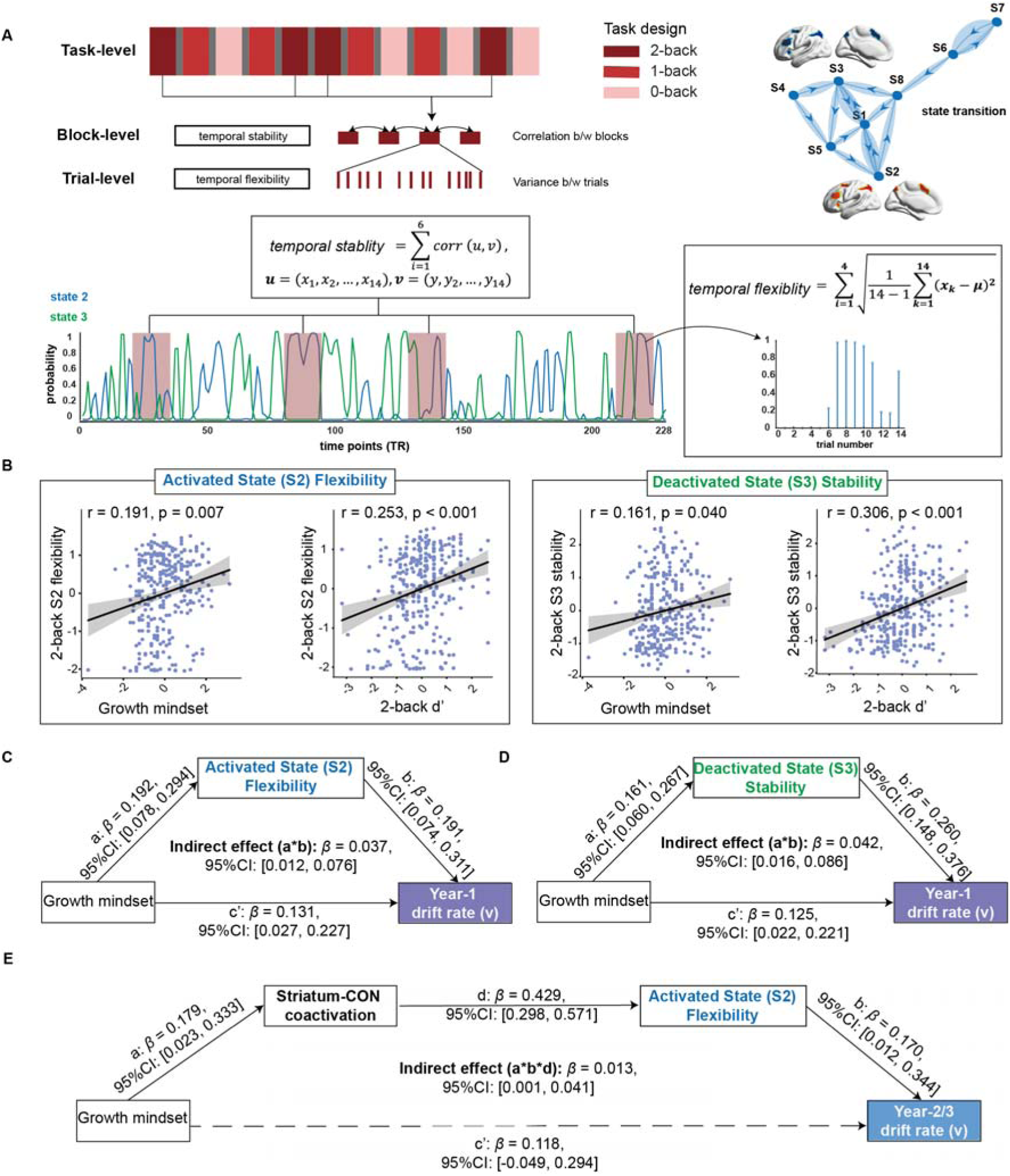
State flexibility and stability of cortico-striatal network dynamics account for the benefits of growth mindset on both cross-sectional and longitudinal WM improvement. (**A**) An overview of the framework for computing temporal flexibility and stability at trial-, block- and task-levels, respectively. Corresponding formulas are provided to compute temporal flexibility and stability metrics. Blue and green lines represent the probability of States 2 and 3 throughout the entire WM task; Green lines: State 3 probability throughout the WM task. (**B**) Regression plots between behavioral performances and state temporal dynamics. Shading represents 95% CI. Each dot represents one subject’s score on the Year-1 test. (**C, D**) Mediation models of State 2 flexibility and State 3 stability under the promotive effect of growth mindset on 2-back WM drift rate at the Year-1 test. (**E**) Chain mediation models with striatum-CON coactivation and State 2 flexibility as sequential mediators.

Given the above observations on prominent correlations among children’s growth mindset, task-invoked activity, latent dynamic states in cortico-striatal networks, and WM performance, we further tested the potential mediatory pathways among these variables using the SEM approach. This analysis revealed the significance of a chain mediation model showing that children’s growth mindset could account for a higher drift rate under the WM 2-back condition, through task-invoked regional activity of striatum-CON systems and the temporal flexibility of cortico-striatal network dynamics reflecting a high-task demand state (i.e., State 2) (indirect Est. = 0.018, 95% CI = [0.006, 0.040]). This model could readily explain the benefit of a growth mindset on longitudinal improvement in drift rate at Year 2 and Year 3 (indirect Est. = 0.013, 95% CI = [0.001, 0.041]). Together, multiple results from our analyses indicate that task-invoked regional activity and nuanced cortico-striatal network dynamics, with flexibility and stability, work in concert to account for the benefits of children’s growth mindset on latent evidence accumulation during information updating in WM.

## Discussion

By integrating pediatric neuroimaging, computational modeling, and brain network dynamics with longitudinal behavioral trajectories, we investigated the neurocognitive mechanisms underlying growth mindset-fostered WM development in youths. Behaviorally, children with higher growth mindset exhibited both cross-sectional and longitudinal improvements in WM performance over three years, along with faster evidence accumulation during moment-to-moment information updating, with the most prominent effect in high task-demand condition. Neurally, growth mindset was associated with greater activation of the dorsal striatum, cingulo-opercular (core part of action-mode network), and fronto-parietal networks during WM, which accounted for developmental improvement in WM performance and latent evidence accumulation. Such activation effects are linked to motivation and cognitive control, and covary with dopaminergic mediators. Further network dynamic modeling revealed that growth mindset optimizes cortico-striatal network dynamics during high task demands, with higher flexibility of an activated state to support trial-by-trial information updating and more stability of a deactivated state across blocks to maintain task goals. Such spatiotemporal features may reflect a balanced and efficient allocation of neurocognitive resources to support changing demands. Our findings underline a neurocognitive framework in which growth mindset enhances latent dynamic computations and effective processing via nuanced orchestration of cortico-striatal networks and fosters far-reaching cognitive development.

### Behavioral and computational processes of growth mindset-fostered WM development

Behaviorally, we observed the benefits of a growth mindset on cross-sectional and longitudinal improvement in WM performance – that is, better accuracy and faster RTs – from middle-to-late childhood into adolescence, with the most prominent effect in the 2-back condition. Our cross-sectional results are in line with previous findings regarding growth mindset benefits on math learning and cognitive control abilities, due to the optimization of cognitive processes(*1*, *14*). Critically, our observed longitudinal effect was consistent across different age groups at baseline (Year-1) and remained robustly evident spanning over three years in Years-2 and 3. Such longitudinal results extend findings from previous cross-sectional studies focusing on growth mindset-fostered effects on behavioral performance in a variety of cognitive tasks, (*3*, *13*, *20*, *25*). Our cross-sectional and longitudinal findings concur with key predictions from influential theories of growth mindset pertaining to motivation-cognition interactions.

Importantly, our observed growth mindset-fostered improvements are attributed to faster speed of evidence accumulation (i.e., drift rate) rather than other latent processes such as decision threshold during information updating, especially under the 2-back condition. Based on the HDDM’s theoretical framework(*26–28*), this finding suggests an enhancement in the efficient processing of latent dynamic computations involved in WM. When performing the 2-back task, one must constantly update and maintain the most recent 2 items in mind and accumulate sufficient evidence extracted from rapidly presented stimuli to ensure a correct decision whether the current item is a target or not(*28*, *51*). In principle, children with growth mindset exhibit core cognitive features centered on malleability beliefs and adaptive processing driven by internal-controllable attributions(*10*, *20*, *52*, *53*). Such traits reflect a dynamic cognitive schema prioritizing task goals and optimizing cognitive-motivational outcomes(*11*, *52*, *54*). This can enhance efficient extraction and accumulation of target-relevant information (evidence) while suppressing irrelevant ones to reach a decision, thereby improving the speed of evidence accumulation. While modulations on lower-level perceptual processes may also occur, our observed benefits are primarily linked to higher-order evidence accumulation of dynamic computations that facilitate efficient processing. Thus, our findings provide novel insights into the computational mechanism of growth mindset-fostered developmental improvements in WM performance.

### Growth mindset fosters WM development through the motivation-cognition dual systems

On the neural level, growth mindset was associated with higher WM-related activation in cortico-striatal networks, including the dorsal striatum, cingulo-opercular, and fronto-parietal systems that covary with dopaminergic modulators. These results are compatible with previous findings on growth mindset-related brain systems localized in the ACC, anterior insula(*11*, *13*) , and striatum(*6*, *31*). This further supports our hypothesis that growth mindset optimizes WM processing by greater engagement of prefrontal-parietal and motivation-related striatal systems(*36*, *55*). Indeed, results from our meta-analytic decoding affirm that the coactivation map (including CON regions) of our observed dorsal striatum is associated with both cognitive and motivational processes, supporting the dual system model of motivation-cognition interplay(*41*, *56*, *57*). Growth mindset-related brain map showed a stronger correlation with attention, WM, and mental computations (i.e., arithmetic, calculation, subtraction), which echoes motivational modulations of growth mindset on attentional control and goal-directed processes(*3*, *11*, *13*). Thus, it is well possible that our observed greater engagement in cortico-striatal systems reflects growth mindset-fostered motivation-cognition interplay. The involvement of motivation-cognition dual systems is also supported by results from our meta-neurotransmitter association analysis. That is, gowth mindset-induced greater activation in cortico-striatal systems covariates with a set of dopaminergic projections and related mediators. The dopaminergic modulations are crucial for goal-directed behaviors via enhancing engagement of cortico-striatal systems, along with motivated efforts and intrinsic evaluation of input information(*58*, *59*). Other neurotransmitters and receptors are also linked to growth mindset-related activation map in our study, including GABA, glutamate, serotonin, and norepinephrine. The balance between glutamate and GABA plays a critical role in excitatory and inhibitory dopaminergic signaling in cortical regions, and modulators like serotonin and norepinephrine facilitate regulating cognitive flexibility(*44*, *60*, *61*).

According to the Self-Determination Theory and neurocognitive models(*18*, *19*, *23*, *24*), our observed greater activation in the dorsal striatum and core nodes of the CON (ACC and anterior insular) may reflect an increase in children’s motivated efforts and internal-controllable values to perform the ongoing WM task, resulting in greater engagement of WM-related prefrontal-parietal systems(*35*, *36*, *41*). Moreover, growth mindset-related cortico-striatal coactivation, especially the striatum-CON, positively correlated with both WM d-prime and drift rate even after two years, connecting these neural correlates with longitudinal outcomes in WM performance. Compared to FPN, enhanced engagement of CON and dorsal striatum may serve to initiate and maintain a high-arousal “action mode” of the brain that enables fast updating and flexible reaction to external demands, leading to more efficient information processing(*45*). In line with our findings, greater coactivation of the striatum and CON may lead to more rapid updating of information in the face of challenges and facilitate adaptive behavior(*62*, *63*). Similar neural patterns have also been seen in the process of intrinsic motivation facilitating high performance in WM tasks(*64*, *65*), supporting our hypothesis that growth mindset enhances motivation and leads to higher cognitive efforts in demanding tasks. Such motivation-cognition dual systems can even predict longitudinal WM outcomes over development from middle childhood to adolescence.

### Growth mindset fosters WM development via optimizing the dynamic orchestration of cortico-striatal networks

In conjunction with cortico-striatal coactivation, growth mindset optimizes the occupancy and transient transitions among eight latent brain states involved in three WM-load conditions. Critically, we identified two brain states with opposite spatiotemporal (re)configuration patterns, characterized by an overall activation (S2) and a deactivation (S3) among core regions of cortico-striatal networks during WM. The S2 occurs more frequently during high task demand (i.e., 2-back), with higher engagement of the dorsal striatum, CON, and FPN regions. As discussed above, higher engagement of these regions assists with motivation-cognition interaction and allows for monitoring and updating of goal-related stimuli(*35*, *36*, *41*). These regions are also linked to cognitive control over information updating, supporting stimulus-triggered transient responses while maintaining relevant information(*59*, *66*). This interpretion is further supported by its positive correlation with faster evidence accumulation. In contrast, the S3 showed higher frequency in low task demands, during which cortico-striatal regions were deactivated in order to reserve cognitive resources(*39*, *67*). Moreover, the frequency of S2 and S3 was positively linked to both growth mindset and WM performance, indicating their crucial roles in growth mindset-fostered WM processing.

Beyond state frequency, our observed associations and mediation pathways among growth mindset and spatiotemporal features of S2 and S3 further underline an intricate interplay of motivational, executive, and action-mode systems anchored onto large-scale cortico-striatal networks. Specifically, the promotive effect of growth mindset on drift rate was mediated by the temporal stability of S3 across blocks under the same goal at baseline, as well as the flexibility of S2 in response to moment-to-moment information updating specific to the 2-back condition, both cross-sectionally and longitudinally. According to brain network dynamics equited for flexible adaptability to changing demands(*50*, *68–71*), our observed high flexibility on a trial-by-trial level most likely reflects more efficient updating and monitoring of perceptual inputs during S2, while the stable S3 is responsible to maintain given task goals and guarantee a stable supply of cognitive resources across blocks of the same task. By integrating temporal flexibility and stability of these states, such features enable a balanced organization of cortico-striatal networks to facilitate flexible allocation of mental resources in response to moment-by-moment information updating, while maintaining the task goal constantly for optimal outcomes. Indeed, our chain mediation results further support the intricate interplay of multiple networks pertaining to the above interpretation: greater task-invoked activity in striatum-CON systems that work in concert with transient state (S2) flexibility rather than deactivated state (S3) stability during WM accounted for growth mindset-fostered developmental improvements in drift rate over three years. Taken together, results from (co)activation, meta-analytic decoding, and network dynamic modeling converge that growth mindset fosters WM development through the intricate interplay of motivational, executive, and action-mode networks that allow for optimizing flexible allocation of neurocognitive resources in response to changing task demands.

### Several limitations

should be noted when interpreting our findings. First, though we recruited children from typical schools, many other variables such as motivation, intelligence, and general cognitive capabilities, should be taken into account in future studies. Second, we have identified brain systems and networks involved in WM linking to growth mindset, but more sophisticated designs with continuous neuroimaging measures and multiple task domains are required to address how growth mindset affects children’s brain, cognitive affective development more broadly. Third, future studies with innovative imaging techniques measuring neural and metabolic activities are required to address the neuromodulatory mechanisms underlying growth mindset-fostered cognitive development.

## In conclusion

our study demonstrates the pivotal role of nuanced cortico-striatal network dynamics with distinct state flexibility and stability in mediating growth mindset-fostered improvements in WM over development in youths. Endorsing a growth mindset appears to optimize the intricate interplay of motivational, executive, and action-mode networks that promote latent dynamic computations and cognitive outcomes. Our findings suggest a neurocognitive account for how growth mindset fosters cognitive development via cortical-striatal networks, which informs interventions for promoting learning programs in education.

## Methods and materials

### Participants

A total of 748 measurements in 454 school-aged children (ranging from 8 to 15 years old) were included in this study, which was derived from the Children School functions and Brain Development Project (CBD, Beijing Cohort). At their first visit, 375 children (mean age ± SD = 9.833±1.039) underwent fMRI scanning while they were performing the N-back task and completed the growth mindset assessment. The average interval between MRI scan and growth mindset assessment was around two months (63 ± 59 days). Children with excessive head motions (more than 1/3 frames with standardized DVARS >1.5 or frame displacement > 0.5) or incomplete scanning were excluded from further analyses **(Figure S1)**. A final sample of 306 children was included in the brain imaging analysis.

In the follow-up test, a subsample of 229 and 144 children were invited back to perform the WM n-back task in the second and third year, respectively **(Figure 1A and 1D)**. To investigate the neural substrates underlying the longitudinal effect of growth mindset on WM development, only children who had high-quality fMRI data at their first visit, along with completed follow-up WM tests, were considered. If children were retested in both the second and third years, the latter observation was selected to represent the longitudinal WM performance with a longer time lag, which resulted in 153 children for the longitudinal subset. Demographic information is summarized in **Table S1**. The written informed consent form was obtained from each child participant and their caregivers or legal guardians. The study procedures were approved by local ethics following the standards of the Declaration of Helsinki. Participants had no obstacle in vision and reported no history of neurological or psychiatric disorders and no current use of any medication or recreational drugs.

### Growth mindset assessment

The Growth Mindset Scale (GMS) (adapted from *15*) consists of 20 items (e.g., No matter who you are, you always can change your intelligence a lot), with 14 items about the individuals’ theory of ability and 6 items about the individuals’ theory of personality (Chinese version used in *5*, *72*). Participants were asked to rate their agreement with each statement using a 4-point Likert-type scale (0 = Strongly Disagree, 1 = Disagree, 2 = Agree, 3 = Strongly Agree). Children aged 9 years old and above completed the questionnaire independently, and children under 9 completed the questionnaire with their parental or assessor’s assistance. The final scores range from 0 to 60, with higher scores representing a higher growth mindset level.

### N-back WM task

A classic numerical N-back task was used to assess participants’ WM performance **(Figure 1B)**. This task consisted of three conditions with three different workloads (i.e., 0-back, 1-back, and 2-back), and each condition consisted of 4 blocks. In each block, participants first viewed a 2-second cue that indicated the workload of this block (i.e., 0-back, 1-back, and 2-back), followed by a sequence of 15 pseudorandom digits in which each digit was presented for 400 milliseconds. In the 0-back condition, participants were instructed to judge whether the current item on the screen was “1” or not by a button press. In the 1-back condition, participants were asked to judge whether the current item was just the same as the previous one. In the 2-back condition, participants needed to judge whether the current item was the same as the one at two positions back. Stimuli were presented via E-Prime 2.0 (http://www.pstnet.com; Psychology Software Tools, Inc., Pittsburgh, PA). Both participants’ response and reaction times (RTs) were recorded. We computed participants’ behavioral performance based on their responses.

### Behavioral performance

The behavioral performance was assessed by the discrimination ability of d-prime (d’) based on the signal detection theory (*73*). All trials for each participant were assigned into the following categories: (1) hits, responses to targets; (2) misses, no response to targets; (3) false alarms, responses to non-targets; (4) correct rejections, no response to non-targets. The hit rate and false alarm rate were defined as follows:

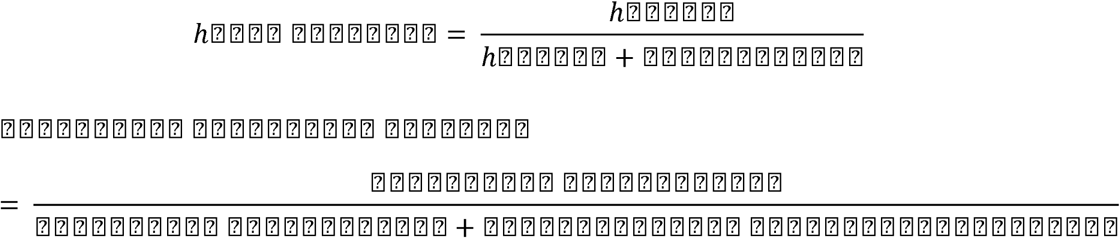

The aforementioned hit rate and false alarm rate were *Z* transformed with inversed cumulative Gaussian distribution to calculate d’ (Finc et al., 2020):

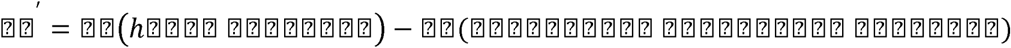

To get finite *d’* values in case either the hit rate or false alarm rate was equal to 0 or 1, modified values 0.01 or 0.99 were used instead. This *d’* measurement was used to assess participants’ WM performance together with RT in all analyses.

### Behavioral statistical analysis

Behavioral measurements were analyzed with R (version 4.0.2, https://www.r-project.org, Platform: x86_64-apple-darwin17.0 (64-bit)). We used the mixed-effect model to evaluate the effect of growth mindset across three years using the lme4 package in R (*74*). The mixed-effect model allows us to test the fixed effects (group-level effect across longitudinal repeats or years in the current study) while controlling for random effects that vary across participants. We first compared models with age and gender as predictors to validate the developmental trajectory of each behavioral measurement. We then constructed a null model with only a fixed intercept and a random intercept to account for the repeated measurements. We further compared the null model with a gender model, a linear age effect model, and a quadratic age effect model. After determining the developmental trajectory for each behavioral measurement, we tested the effects of growth mindset on behavioral performance over age and gender. Models were fitted with the maximum likelihood (ML) estimation method. Model comparison was based on both Akaike Information Criterion (AIC) and Bayesian information criterion (BIC).

### Mediation analysis

Before mediation analysis, we calculated the correlation coefficients of each ROI activation with mindset levels, WM performances, and HDDM parameters. After correcting for multiple comparisons using FDR, networks with at least one region related to mindset and behavioral outcomes were sent for further mediation analysis. We conducted and statistically tested all mediation effects using Mplus 8.3 (*75*). Firstly, the structural equation models were constructed to examine whether the overall FPN and striatum-CON coactivation mediated the influence of growth mindset on current and future working memory performance separately. Based on the results from connectivity analysis, we constructed specific models to investigate how growth mindset influences current and future latent decision-making processes (drift rate) through cortico-striatal interaction and regional response in the insula. Because the effect of age and gender had already been controlled in each analysis of activation and connectivity, these two covariates were included in the equation of behavioral outcomes. The fitness of each model was assessed using a _χ_2 test, which showed no significance. In addition, the Root Mean Square Error of Approximation (RMSEA) was found to be below 0.08, the Standardized Root Mean Square Residual (SRMR) had an outcome value of less than 0.08, and the Comparative Fit Index (CFI) was above 0.90. For all models, a number of 5000 draws were done for bootstrap, and 95% confidence intervals were estimated. If the confidence interval did not include zero, the effect was considered significant.

### HDDM for N-back task

The DDM describes decision-making as an evidence-accumulation process in which individuals continually accumulate effective evidence from the environment until they reach an internal threshold to make a final decision. Based on trail-by-trail RTs and accuracies, the DDM decomposes participants’ decision-making performances into latent processes, which were indicated by three free parameters: drift rate (v), decision threshold (a), and non-decision time (t) (*76*). It is widely utilized in two-choice decision tasks but is also extendedly implemented for one-choice tasks such as Go/no-Go tasks (*27*, *77*) and N-back tasks (*30*). In such paradigms, only trail-by-trail RTs for hits and false alarms were fitted for the DDM, while RTs of misses and correct rejections cannot be measured. For the current N-back task, the latent process is represented by participants accumulating evidence during a series of digit stimulus presentations to decide whether the current digit is the same as the digits that appear one or two points before in the sequence and then make the corresponding response. We estimated the DDM parameters by HDDM because it is more applicable to the case of relatively few trials and is able to estimate both group-level and individual-level parameters by drawing individual parameters from the group distribution. In this way, we can obtain group- and individual-level parameters for three workloads at three time points under the current paradigm. Note that models were estimated using all data with test timepoint and workloads were set as factors.

To exclude the effect of different incorporated parameters on the results, we fitted seven models with the same data, including models of a, v, t, av, at, vt, avt. For each model, Markov chain Monte Carlo (MCMC) sampling method was applied to generate 20,000 samples and discard the initial 2,000 samples as burn-in for performing Bayesian inference. We then compared the models based on the deviance information criterion (DIC) (*78*). The model with the lowest DIC value was considered to have the best fit. Gelman-Rubin statistics were further employed to assess the convergence of the model. The r-hat values for each parameter were close to 1.0 and less than 1.1, indicating good convergence (*76*). Finally, the three individual-level parameters of the best-fit model (avt) were submitted to subsequent analyses (**Figure S5 & Table S2**).

### Image data acquisition

Data were acquired using the same type of 3.0T scanner (Magnetom Prisma syngo MR D13D, Erlangen, Germany) with a 64-channel head coil from two sites. High-resolution anatomical images were acquired by a three-dimensional sagittal T1-weighted magnetization-prepared rapid gradient echo (MPRAGE) sequence (TR = 2530 ms, TE = 2.98 ms, TI = 1100ms, flip angle = 7°, voxel size 0.5 x 0.5 x 1.0 mm3, matrix size = 256 × 224, FOV = 256 x 224 mm2, brand width = 240 Hz/Px, 192 slices with 1 mm thickness). Functional images with 33 axial slices (3.5 mm thick, 0.7 mm skip) parallel to the anterior and posterior commissural line (AC-PC) were acquired using a T2*-sensitive echo-planar imaging (EPI) sequence (TR = 2000ms, TE = 30ms, flip angle = 90°, voxel size = 3.5 x 3.5 x 3.5 mm3, FOV = 224 x 224 mm2). The whole n-back WM task consisted of 232 volumes and lasted for 464s. Two different sites were included as a nuisance for fMRI data analyses.

### Image data preprocessing

Brain images were preprocessed with the fMRIPrep 1.4.1 (*79*) pipeline implemented in Nipype 1.2.0 (*80*). The first 4 volumes of n-back task were discarded for signal stability and adaptation of participants. For each participant, the following preprocessing procedures were conducted. First, each T1w volume was skull-stripped and a BOLD reference was estimated. Slice time correction was then performed and all slices were realigned in time to the middle of each TR using 3dTshift from AFNI. Motion correction was done using mcflirt (FSL) and 6 head-motion parameters (three rotations, three translations) were estimated. The EPI data was corrected for susceptibility distortions based on a field map and co-registered to the anatomical reference using boundary-based registration with nine degrees of freedom. Finally, these preprocessed BOLD functional images in the original space were resampled into the well-known ‘MNI152NLin6Asym’ space. Head-motion transformation, susceptibility distortion correction, BOLD-to-T1w transformation and T1w-to-template (MNI) warp were concatenated and applied in a single step using antsApplyTransforms (ANTs) using Lanczos interpolation.

ICA-based Automatic Removal of Motion Artifacts (ICA-AROMA) was used to automatically remove motion artifacts non-aggressively after removal of non-steady volumes and spatial smoothing with an isotropic, Gaussian kernel of 6 mm full-width half-maximum (FWHM). Physiological noise regressors were extracted applying CompCor and two CompCor variants were estimated: temporal (tCompCor) and anatomical (aCompCor). Framewise displacement (FD) and DVARS were calculated using Nipype. In addition to 6 head-motion parameters and global signals, their temporal derivatives and quadratic terms were also estimated. Outliers were defined as frames that exceeded a threshold of 0.5 mm FD and 1.5 standardized DVARS and were annotated. All these parameters were taken as aggressive noise regressors and were placed in the corresponding confounds file. For credible results, individuals with more than 1/3 frames as outliers were excluded for further analyses (n = 22).

### General linear model (GLM) analysis

To identify WM-related brain systems and their relations to growth mindset, we constructed GLMs on both individual and group levels using SPM12 (https://www.fil.ion.ucl.ac.uk/spm/software/spm12/). To assess task-invoked neural response to different workloads including 0-, 1-, and 2-back conditions were modeled as separate boxcar regressors and convolved with the canonical hemodynamic response function (HRF) built in SPM12. To regress out effects related to noise, signals within cerebrospinal fluid and white matter from each participant were included as a nuisance in the model (Parkes et al., 2018). A high-pass filter of 1/128Hz was applied and temporal autocorrelations in fMRI were corrected using a first-order autoregressive model (AR(1)).

Relevant contrast parameter estimate images were initially generated at the individual-subject level, and then submitted to group-level analyses by treating participants as a random variable. Contrast images of 2-back > 0-back, 2-back > 1-back, and 1-back > 0-back were submitted to separate multiple regression analyses with children’s growth mindset as a covariate of interest, gender and age as nuisances. Coefficients of the multiple regression maps were tested using one-sample t-test. Significant clusters were determined at a voxel-level false discovery rate (FDR) correction (pFDR < 0.05) on the whole brain. For visualization of results, significant clusters were displayed using Surf Ice (https://www.nitrc.org/projects/surfice/) and MRIcroGL (https://www.nitrc.org/projects/mricrogl/).

### Meta-analytic coactivation and decoding

The meta-analytic coactivation map is a kind of meta-analog of functional connectivity map and is generated by the Neurosynth platform (http://neurosynth.org) based on the coordinates of seed regions. Our analysis is seeded in a 6-mm sphere centered on the coordinates of peak activity in the bilateral caudate nucleus identified by the multiple regression analysis for WM-related brain activity (2-versus 0-back contrast) with the growth mindset as a covariate of interest. The map reflects coactivation of brain regions across studies in the Neurosynth database and voxels with high Z values in the map are likely to be activated in similar studies as the seed voxels. The final coactivation mask only included clusters with more than 10 voxels satisfied Z > 3 (*p* < 0.001) in coactivation maps of both the left and right caudate nuclei. Subsequently, we overlapped the coactivation mask with the multiple regression results of growth mindset for comparison and visualization purposes. Mask of multiple comparisons only included continuous clusters with more than 10 voxels passed the threshold of p < 0.05 corrected for multiple comparisons using FDR in the 2-back > 0-back contrast.

We then uploaded these two masks to the Image Decoder based on the Neurosynth database to identify the most related terms of brain response. The top 10 functional terms whose neural response showed the highest correlation with the current mask were presented. Because five terms were decoded in both masks, the final result presented 15 terms in total. Note that the redundant or anatomical terms were not included (e.g., “working memory” was presented, and “working” or “caudate” were removed).

### Neurotransmitter Analysis

To investigate the neurochemical characteristics of the identified growth mindset, we calculated its spatial similarity with a series of PET tracer neurotransmitter receptor/transporter maps (*81*). This PET tracer dataset is independent of this study and includes mean neurochemical binding maps from 36 studies with over 1200 healthy individuals. These studies provide independent maps for 19 receptors and transporters across nine neurotransmitter systems, including cannabinoids, dopamine, norepinephrine, acetylcholine, serotonin, GABA, histamine, glutamate, and opioids, which were then resampled to standard space for assessing spatial similarity.

We first ran 100 times bootstrapping of the multiple regression described above. In each bootstrapping iteration, participants’ 2-back > 0-back contrast map was resampled and then used to create a growth mindset map through multiple regression. After resampling the neurotransmitter system into imaging space, we calculated the voxel-by-voxel Pearson correlation between the bootstrapped growth mindset map and each neurotransmitter receptor/transporter map to estimate their spatial similarity. For all bootstrap samples, we computed the mean spatial correlations and the corresponding estimated standard error (the standard deviation), which were both presented in the Figure 2c radar plot. We only interpreted neurochemical similarities found in neurotransmitters validated across studies, meaning those with 95% confidence intervals above zero in individual PET studies. The bootstrapping method and replications across studies ensured the validation of our findings.

### Regions of interest (ROIs) analysis

ROIs of three functional brain networks were generated from meta-analysis images using the Neurosynth database. We searched for the meta-analysis maps related to terms “working memory” and “default mode”, and converted the maps to masks only including clusters with more than 10 voxels satisfied Z > 3 (p < 0.001). The whole-brain result and meta-analysis results were also overlapped for comparison and visualization.

Based on this criterion, we then picked the caudate nucleus for the striatum, anterior Insula (aIns), and dorsal anterior cingulate cortex (dACC) for the CON from the meta-analysis map of the term “working memory”. These regions in the striatum-CON system were responsible for motivational response during cognitive tasks as aforementioned. For the FPN, dorsal lateral prefrontal cortex (dlPFC), frontal eye field (FEF), and inferior parietal sulcus (IPS) were selected from the same map. Likewise, clusters of angular gyrus (AG), posterior cingulate cortex (PCC), and ventral medial prefrontal cortex (vmPFC) were selected from the meta-analysis map of the term “default mode”. Besides, the bilateral hippocampus/parahippocampus (HCP/PHC) was also included based on the result of whole-brain multiple regression. Finally, ROIs generated from the original meta-analysis maps include: caudate for striatum; aIns and dACC for CON; dlPFC, FEF, and IPS for FPN; vmPFC, AG, PCC, and HCP/PHC for DMN **(Figure 3A & S6)**. Parameter estimates from each ROI, and each participant was extracted from the individual-level contrast of 2-back with the 0-back condition using the MarsBaR (http://marsbar.sourceforge.net/) to characterize activation during the task in each ROI.

### Hidden Markov Model (HMM)

We implemented the HMM to model the spatiotemporal brain dynamics within the mesolimbic and frontoparietal systems with which children meet the changing requirements and demands during the WM task. The HMM captured a set of distinct brain states from the multidimensional input signals extracted from 11 ROIs within striatum-CON and FPN. These brain states were determined using Gaussian distribution, taking into account the mean of each dimension and covariance among dimensions. The HMM-MAR toolbox (https://github.com/OHBA-analysis/HMM-MAR) was utilized to estimate the changing brain configurations underlying the data. To decide on the final number of states, we compared the models of 6, 8, 10, and 12 states and selected the model whose output best aligned with our predefined workload changes. Through model comparison, we confirmed that the temporal features of 8-state outputs are sufficient to cover the neural dynamics of three workloads, while the others with higher than 8 states have inefficient states that were rarely active throughout the whole task. After deciding the final number of states, the Viterbi algorithm was used to obtain the maximum a posteriori probability path. As shown in **Figure 4**, the final temporospatial outputs included the decoded mostly likely sequence of hidden states, the frequency of occurrence for each state under each condition, the transition metrics among the 8 states, and the spatial features of each state within the predefined brain system. The temporospatial features were then correlated with behavioral indices to select our interested states for further analysis, and the occurrence probability was used to calculate the temporal stability and flexibility of each state sequence.

## References

1. D. Yeager, P. Hanselman, G. Walton, J. Murray, R. Crosnoe, C. Muller, E. Tipton, B. Schneider, C. Hulleman, C. Hinojosa, et al., A national experiment reveals where a growth mindset improves achievement. Nature 573, 364–369 (2019). 10.1038/s41586-019-1466-y.

2. C. Dweck, D. Yeager, Mindsets: A View From Two Eras. Perspect. Psychol. Sci. 14 (2019). 10.1177/1745691618804166.

3. H. S. Schroder, M. E. Fisher, Y. Lin, S. L. Lo, J. H. Danovitch, J. S. Moser, Neural evidence for enhanced attention to mistakes among school-aged children with a growth mindset. Dev. Cogn. Neurosci. 24, 42–50 (2017). 10.1016/j.dcn.2017.01.004.

4. J. B. Sarrasin, L. Nenciovici, L. M. B. Foisy, G. Allaire-Duquette, M. Riopel, S. Masson, Effects of teaching the concept of neuroplasticity to induce a growth mindset on motivation, achievement, and brain activity: A meta-analysis. Elsevier GmbH (2018). 10.1016/j.tine.2018.07.003.

5. D. Wang, L. Gan, C. Wang, The effect of growth mindset on reasoning ability in Chinese adolescents and young adults: the moderating role of self-esteem. Curr. Psychol., 1–7 (2021). 10.1007/s12144-021-01437-9.

6. L. Chen, H. Chang, J. Rudoler, E. Arnardottir, Y. Zhang, C. de los Angeles, V. Menon, Cognitive training enhances growth mindset in children through plasticity of cortico-striatal circuits. *npj Sci*. Learn. 7, 1–10 (2022). 10.1038/s41539-022-00146-7.

7. R. A. I. Bethlehem, J. Seidlitz, S. R. White, J. W. Vogel, K. M. Anderson, C. Adamson, S. Adler, G. S. Alexopoulos, E. Anagnostou, A. Areces-Gonzalez, et al., Brain charts for the human lifespan. Nature 604, 525–533 (2022). 10.1038/s41586-022-04554-y.

8. B. Tervo-Clemmens, F. J. Calabro, A. C. Parr, J. Fedor, W. Foran, B. Luna, A Canonical Trajectory of Executive Function Maturation During the Transition from Adolescence to Adulthood. Nat. Commun. 14, 1–17 (2023). 10.1038/s41467-023-42540-8.

9. K. Ganesan, A. Thompson, C. R. Smid, R. Cañigueral, Y. Li, G. Revill, V. Puetz, B. C. Bernhardt, N. U. F. Dosenbach, R. Kievit, et al., Cognitive control training with domain-general response inhibition does not change children’s brains or behavior. Nat. Neurosci. 2024 277 27, 1364–1375 (2024). 10.1038/s41593-024-01672-w.

10. D. Yeager, C. Bryan, J. Gross, D. Krettek, P. Santos, J. Murray, H. Graveling, M. Johnson, J. Jamieson, A Synergistic Mindsets Intervention Protects Adolescents from Social Stress. Nature 607, 512–520 (2021). 10.21203/rs.3.rs-551170/v1.

11. J. Mangels, B. Butterfield, J. Lamb, C. Good, C. Dweck, Why do beliefs about intelligence influence learning success? A social cognitive neuroscience model. Soc. Cogn. Affect. Neurosci. 1, 75–86 (2006). 10.1093/scan/nsl013.

12. H. S. Schroder, T. P. Moran, M. B. Donnellan, J. S. Moser, Mindset induction effects on cognitive control: A neurobehavioral investigation. Biol. Psychol. 103, 27–37 (2014). 10.1016/J.BIOPSYCHO.2014.08.004.

13. J. S. Moser, H. S. Schroder, C. Heeter, T. P. Moran, Y. H. Lee, Mind your errors: Evidence for a neural mechanism linking growth mind-set to adaptive posterror adjustments. Psychol. Sci. 22, 1484–1489 (2011). 10.1177/0956797611419520.

14. L. Blackwell, K. Trzesniewski, C. Dweck, Implicit theories of intelligence predict achievement across an adolescent transition: A longitudinal study and an intervention. Child Dev. 78, 246–263 (2007). 10.1111/j.1467-8624.2007.00995.x.

15. C. Dweck, Mindset: The New Psychology of Success. (Random House, New York, NY, US, 2006).

16. F. Nemmi, C. Nymberg, E. Helander, T. Klingberg, Grit Is Associated with Structure of Nucleus Accumbens and Gains in Cognitive Training. J. Cogn. Neurosci. 28, 1688– 1699 (2016). 10.1162/JOCN_A_01031.

17. T. Salminen, S. Kühn, P. A. Frensch, T. Schubert, Transfer after dual n-back training depends on striatal activation change. J. Neurosci. 36, 10198–10213 (2016). 10.1523/JNEUROSCI.2305-15.2016.

18. R. M. Ryan, E. L. Deci, Self-determination theory and the facilitation of intrinsic motivation, social development, and well-being. Am. Psychol. 55, 68–78 (2000). 10.1037/0003-066X.55.1.68.

19. R. M. Ryan, E. L. Deci, Intrinsic and extrinsic motivation from a self-determination theory perspective: Definitions, theory, practices, and future directions. Contemp. Educ. Psychol. 61, 101860 (2020). 10.1016/J.CEDPSYCH.2020.101860.

20. K. M. Xu, P. Koorn, B. de Koning, I. T. Skuballa, L. Lin, M. Henderikx, H. W. Marsh, J. Sweller, F. Paas, A Growth Mindset Lowers Perceived Cognitive Load and Improves Learning: Integrating Motivation to Cognitive Load. J. Educ. Psychol., doi: 10.1037/edu0000631 (2020).

21. C. Ames, J. Archer, Achievement Goals in the Classroom: Students’ Learning Strategies and Motivation Processes. J. Educ. Psychol. 80, 260–267 (1988).

22. C. Dweck, “Self-theories motivate self-regulated learning” in Motivation and Self-Regulated Learning (Routledge, 2012), pp. 31–51.

23. R. J. Spiro, R. L. Coulson, P. J. Feltovich, D. K. Anderson, Cognitive Flexibility Theory: Advanced Knowledge Acquisition in Ill-Structured Domains. Proc. Annu. Meet. Cogn. Sci. Soc. 10, 0 (1988).

24. L. Q. Uddin, Cognitive and behavioural flexibility: neural mechanisms and clinical considerations. Nat. Rev. Neurosci. 2021 223 22, 167–179 (2021). 10.1038/s41583-021-00428-w.

25. H. S. Schroder, T. P. Moran, M. B. Donnellan, J. S. Moser, Mindset induction effects on cognitive control: A neurobehavioral investigation. Biol. Psychol. 103, 27–37 (2014). 10.1016/J.BIOPSYCHO.2014.08.004.

26. R. Ratcliff, G. McKoon, The diffusion decision model: Theory and data for two-choice decision tasks. Neural Comput. 20, 873–922 (2008). 10.1162/NECO.2008.12-06-420.

27. R. Ratcliff, C. Huang-Pollock, G. McKoon, Modeling individual differences in the go/no-go task with a diffusion model. Decision 5, 42–62 (2018). 10.1037/DEC0000065.

28. M. L. Pedersen, D. Alnæs, D. van der Meer, S. Fernandez-Cabello, P. Berthet, A. Dahl, R. Kjelkenes, E. Schwarz, W. K. Thompson, D. M. Barch, et al., Computational Modeling of the n-Back Task in the ABCD Study: Associations of Drift Diffusion Model Parameters to Polygenic Scores of Mental Disorders and Cardiometabolic Diseases. Biol. Psychiatry Cogn. Neurosci. Neuroimaging 8, 290–299 (2023). 10.1016/j.bpsc.2022.03.012.

29. R. Ratcliff, P. L. Smith, S. D. Brown, G. McKoon, Diffusion Decision Model: Current Issues and History. Trends Cogn. Sci. 20, 260–281 (2016). 10.1016/J.TICS.2016.01.007.

30. L. Liu, J. Wu, H. Geng, C. Liu, Y. Luo, J. Luo, S. Qin, Long-term stress and trait anxiety affect brain network balance in dynamic cognitive computations. Cereb. Cortex 32, 2957–2971 (2022). 10.1093/cercor/bhab393.

31. C. A. Myers, C. Wang, J. M. Black, N. Bugescu, F. Hoeft, The matter of motivation: Striatal resting-state connectivity is dissociable between grit and growth mindset. Soc. Cogn. Affect. Neurosci. 11, 1521–1527 (2016). 10.1093/scan/nsw065.

32. S. Palminteri, D. Justo, C. Jauffret, B. Pavlicek, A. Dauta, C. Delmaire, V. Czernecki, C. Karachi, L. Capelle, A. Durr, et al., Critical Roles for Anterior Insula and Dorsal Striatum in Punishment-Based Avoidance Learning. Neuron 76, 998–1009 (2012). 10.1016/J.NEURON.2012.10.017.

33. C. H. Chatham, M. J. Frank, D. Badre, Corticostriatal output gating during selection from working memory. Neuron 81, 930–942 (2014). 10.1016/j.neuron.2014.01.002.

34. D. G. Ghahremani, B. Lee, C. L. Robertson, G. Tabibnia, A. T. Morgan, N. De Shetler, A. K. Brown, J. R. Monterosso, A. R. Aron, M. A. Mandelkern, et al., Striatal Dopamine D2/D3 Receptors Mediate Response Inhibition and Related Activity in Frontostriatal Neural Circuitry in Humans. J. Neurosci. 32, 7316–7324 (2012). 10.1523/JNEUROSCI.4284-11.2012.

35. A. Westbrook, R. van den Bosch, J. I. Määttä, L. Hofmans, D. Papadopetraki, R. Cools, M. J. Frank, Dopamine promotes cognitive effort by biasing the benefits versus costs of cognitive work. Science (80-. ). 367, 1362–1366 (2020). 10.1126/science.aaz5891.

36. R. M. Krebs, C. N. Boehler, K. C. Roberts, A. W. Song, M. G. Woldorff, The involvement of the dopaminergic midbrain and cortico-striatal-thalamic circuits in the integration of reward prospect and attentional task demands. Cereb. Cortex 22, 607– 615 (2012). 10.1093/cercor/bhr134.

37. T. Braver, M. Krug, K. Chiew, W. Kool, A. Westbrook, N. Clement, A. Adcock, D. Barch, M. Botvinick, C. Carver, et al., Mechanisms of motivation-cognition interaction: Challenges and opportunities. Springer (2014). 10.3758/s13415-014-0300-0.

38. M. Botvinick, T. Braver, Motivation and cognitive control: From behavior to neural mechanism. Annu. Rev. Psychol. 66, 83–113 (2015). 10.1146/annurev-psych-010814-015044.

39. D. Barulli, Y. Stern, Efficiency, capacity, compensation, maintenance, plasticity: Emerging concepts in cognitive reserve. Trends Cogn. Sci. 17, 502–509 (2013). 10.1016/j.tics.2013.08.012.

40. E. J. Hermans, M. J. A. G. Henckens, M. Joëls, G. Fernández, Dynamic adaptation of large-scale brain networks in response to acute stressors. Trends Neurosci. 37, 304– 314 (2014). 10.1016/J.TINS.2014.03.006.

41. A. R. Arulpragasam, J. A. Cooper, M. R. Nuutinen, M. T. Treadway, Corticoinsular circuits encode subjective value expectation and violation for effortful goal-directed behavior. Proc. Natl. Acad. Sci. U. S. A. 115, E5233–E5242 (2018). 10.1073/pnas.1800444115.

42. P. L. Croxson, M. E. Walton, J. X. O’Reilly, T. E. J. Behrens, M. F. S. Rushworth, Effort-Based Cost–Benefit Valuation and the Human Brain. J. Neurosci. 29, 4531– 4541 (2009). 10.1523/JNEUROSCI.4515-08.2009.

43. K. D’Ardenne, N. Eshel, J. Luka, A. Lenartowicz, L. E. Nystrom, J. D. Cohen, Role of prefrontal cortex and the midbrain dopamine system in working memory updating. Proc. Natl. Acad. Sci. 109, 19900–19909 (2012). 10.1073/PNAS.1116727109.

44. T. Ott, A. Nieder, Dopamine and Cognitive Control in Prefrontal Cortex. Elsevier Current Trends (2019). 10.1016/j.tics.2018.12.006.

45. N. U. F. Dosenbach, M. E. Raichle, E. M. Gordon, The brain’s action-mode network. Nat. Rev. Neurosci., 1–11 (2025). 10.1038/s41583-024-00895-x.

46. M. Ullsperger, C. Danielmeier, G. Jocham, Neurophysiology of performance monitoring and adaptive behavior. Physiol. Rev. 94, 35–79 (2014). 10.1152/physrev.00041.2012.

47. Z. Fu, A. Sajad, S. P. Errington, J. D. Schall, U. Rutishauser, Neurophysiological mechanisms of error monitoring in human and non-human primates. Nature Publishing Group (2023). 10.1038/s41583-022-00670-w.

48. C. Chang, G. H. Glover, Time–frequency dynamics of resting-state brain connectivity measured with fMRI. Neuroimage 50, 81–98 (2010). 10.1016/J.NEUROIMAGE.2009.12.011.

49. R. M. Hutchison, T. Womelsdorf, E. A. Allen, P. A. Bandettini, V. D. Calhoun, M. Corbetta, S. Della Penna, J. H. Duyn, G. H. Glover, J. Gonzalez-Castillo, et al., Dynamic functional connectivity: Promise, issues, and interpretations. Neuroimage 80, 360–378 (2013). 10.1016/j.neuroimage.2013.05.079.

50. Y. Zeng, B. Xiong, H. Gao, C. Liu, C. Chen, J. Wu, S. Qin, Cortisol awakening response prompts dynamic reconfiguration of brain networks in emotional and executive functioning. Proc. Natl. Acad. Sci. 121, e2405850121 (2024). 10.1073/PNAS.2405850121.

51. L. Liu, J. Wu, H. Geng, C. Liu, Y. Luo, J. Luo, S. Qin, Long-term stress and trait anxiety affect brain network balance in dynamic cognitive computations. Cereb. Cortex 32, 2957–2971 (2022). 10.1093/CERCOR/BHAB393.

52. I. Daly, J. Bourgaize, A. Vernitski, Mathematical mindsets increase student motivation: Evidence from the EEG. Trends Neurosci. Educ. 15, 18–28 (2019). 10.1016/J.TINE.2019.02.005.

53. D. S. Yeager, C. S. Dweck, S. Yeager, What Can Be Learned From Growth Mindset Controversies? Association 2020, 1269–1284 (2020). 10.1037/amp0000794.supp.

54. L. Chen, S. R. Bae, C. Battista, S. Qin, T. Chen, T. M. Evans, V. Menon, Positive Attitude Toward Math Supports Early Academic Success: Behavioral Evidence and Neurocognitive Mechanisms. Psychol. Sci. 29, 390–402 (2018). 10.1177/0956797617735528.

55. K. L. Mills, D. Bathula, T. G. C. Dias, S. P. Iyer, M. C. Fenesy, E. D. Musser, C. A. Stevens, B. L. Thurlow, S. D. Carpenter, B. J. Nagel, et al., Altered cortico-striatal-thalamic connectivity in relation to spatial working memory capacity in children with ADHD. Front. Psychiatry 3, 2 (2012). 10.3389/FPSYT.2012.00002/XML/NLM.

56. J. Jiang, J. Beck, K. Heller, T. Egner, An insula-frontostriatal network mediates flexible cognitive control by adaptively predicting changing control demands. Nat. Commun. 6, 1–11 (2015). 10.1038/ncomms9165.

57. D. M. Yee, J. L. Crawford, B. Lamichhane, T. S. Braver, Dorsal Anterior Cingulate Cortex Encodes the Integrated Incentive Motivational Value of Cognitive Task Performance. J. Neurosci. 41, 3707–3720 (2021). 10.1523/JNEUROSCI.2550-20.2021.

58. I. C. Ballard, V. P. Murty, R. M. Carter, J. J. MacInnes, S. A. Huettel, R. A. Adcock, Dorsolateral Prefrontal Cortex Drives Mesolimbic Dopaminergic Regions to Initiate Motivated Behavior. J. Neurosci. 31, 10340–10346 (2011). 10.1523/JNEUROSCI.0895-11.2011.

59. M. Botvinick, T. Braver, Motivation and Cognitive Control: From Behavior to Neural Mechanism. Annu. Rev. Psychol 66, 83–113 (2015). 10.1146/annurev-psych-010814-015044.

60. J. M. Shine, E. J. Müller, B. Munn, J. Cabral, R. J. Moran, M. Breakspear, Computational models link cellular mechanisms of neuromodulation to large-scale neural dynamics. (2021). 10.1038/s41593-021-00824-6.

61. E. Seu, A. Lang, R. J. Rivera, J. D. Jentsch, Inhibition of the norepinephrine transporter improves behavioral flexibility in rats and monkeys. Psychopharmacology (Berl). 202, 505–519 (2009). 10.1007/s00213-008-1250-4.

62. S. Kühn, F. Schmiedek, H. Noack, E. Wenger, N. C. Bodammer, U. Lindenberger, M. Lövden, The dynamics of change in striatal activity following updating training. Hum. Brain Mapp. 34, 1530–1541 (2013). 10.1002/HBM.22007.

63. E. Dahlin, A. S. Neely, A. Larsson, L. Bäckman, L. Nyberg, Transfer of learning after updating training mediated by the striatum. Science (80-. ). 320, 1510–1512 (2008). 10.1126/science.1155466.

64. S. I. Di Domenico, R. M. Ryan, The emerging neuroscience of intrinsic motivation: A new frontier in self-determination research. Frontiers Media S. A (2017). 10.3389/fnhum.2017.00145.

65. T. D. Satterthwaite, K. Ruparel, J. Loughead, M. A. Elliott, R. T. Gerraty, M. E. Calkins, H. Hakonarson, R. C. Gur, R. E. Gur, D. H. Wolf, Being right is its own reward: Load and performance related ventral striatum activation to correct responses during a working memory task in youth. Neuroimage 61, 723–729 (2012). 10.1016/j.neuroimage.2012.03.060.

66. T. S. Braver, The variable nature of cognitive control: A dual mechanisms framework. Trends Cogn. Sci. 16, 106–113 (2012). 10.1016/J.TICS.2011.12.010/ASSET/AE7321A3-40A0-41B8-B78D-2A67C9923B95/MAIN.ASSETS/GR2.JPG.

67. S. Heinzel, R. C. Lorenz, W. R. Brockhaus, T. Wüstenberg, N. Kathmann, A. Heinz, M. A. Rapp, Working memory load-dependent brain response predicts behavioral training gains in older adults. J. Neurosci. 34, 1224–1233 (2014). 10.1523/JNEUROSCI.2463-13.2014.

68. W. Cai, S. Ryali, R. Pasumarthy, V. Talasila, V. Menon, Dynamic causal brain circuits during working memory and their functional controllability. Nat. Commun. 12, 1–16 (2021). 10.1038/s41467-021-23509-x.

69. U. Braun, A. Schäfer, H. Walter, S. Erk, N. Romanczuk-Seiferth, L. Haddad, J. I. Schweiger, O. Grimm, A. Heinz, H. Tost, et al., Dynamic reconfiguration of frontal brain networks during executive cognition in humans. Proc. Natl. Acad. Sci. U. S. A. 112, 11678–11683 (2015). 10.1073/pnas.1422487112.

70. C. V. Cocuzza, T. Ito, D. Schultz, D. S. Bassett, M. W. Cole, Flexible coordinator and switcher hubs for adaptive task control. J. Neurosci. 40, 6949–6968 (2020). 10.1523/JNEUROSCI.2559-19.2020.

71. J. M. Shine, P. G. Bissett, P. T. Bell, O. Koyejo, J. H. Balsters, K. J. Gorgolewski, C. A. Moodie, R. A. Poldrack, The Dynamics of Functional Brain Networks: Integrated Network States during Cognitive Task Performance. Neuron 92, 544–554 (2016). 10.1016/J.NEURON.2016.09.018.

72. D. Wang, F. Yuan, Y. Wang, Growth mindset and academic achievement in Chinese adolescents: A moderated mediation model of reasoning ability and self-affirmation. Curr. Psychol., 1–10 (2020). 10.1007/S12144-019-00597-Z/FIGURES/5.

73 . D. M. Green, J. A. Swets, “SIGNAL DETECTION THEORY AND PSYCHOPHYSICS” (1966).

74. D. Bates, M. Mächler, B. M. Bolker, S. C. Walker, Fitting Linear Mixed-Effects Models Using lme4. J. Stat. Softw. 67, 1–48 (2015). 10.18637/JSS.V067.I01.

75. L. K. Muthén, B. O. Muthén, Mplus user’s guide. Los Angeles. CA Muthén Muthén 2017 (1998).

76. T. V. Wiecki, I. Sofer, M. J. Frank, HDDM: Hierarchical bayesian estimation of the drift-diffusion model in Python. Front. Neuroinform. 7, 14 (2013). 10.3389/FNINF.2013.00014/BIBTEX.

77. J. Zhang, T. Rittman, C. Nombela, A. Fois, I. Coyle-Gilchrist, R. A. Barker, L. E. Hughes, J. B. Rowe, Different decision deficits impair response inhibition in progressive supranuclear palsy and Parkinson’s disease. Brain 139, 161–173 (2016). 10.1093/BRAIN/AWV331.

78. D. J. Spiegelhalter, N. G. Best, B. P. Carlin, A. Van Der Linde, Bayesian measures of model complexity and fit. J. R. Stat. Soc. Ser. B (Statistical Methodol. 64, 583–639 (2002). 10.1111/1467-9868.00353.

79. O. Esteban, C. J. Markiewicz, R. W. Blair, C. A. Moodie, A. I. Isik, A. Erramuzpe, J. D. Kent, M. Goncalves, E. DuPre, M. Snyder, et al., fMRIPrep: a robust preprocessing pipeline for functional MRI. Nat. Methods 16, 111–116 (2019). 10.1038/s41592-018-0235-4.

80. K. Gorgolewski, C. D. Burns, C. Madison, D. Clark, Y. O. Halchenko, M. L. Waskom, S. S. Ghosh, Nipype: A flexible, lightweight and extensible neuroimaging data processing framework in Python. Front. Neuroinform. 5, 13 (2011). 10.3389/fninf.2011.00013.

81. J. Y. Hansen, G. Shafiei, R. D. Markello, K. Smart, S. M. L. Cox, M. Nørgaard, V. Beliveau, Y. Wu, J. D. Gallezot, É. Aumont, et al., Mapping neurotransmitter systems to the structural and functional organization of the human neocortex. Nat. Neurosci. 25, 1569–1581 (2022). 10.1038/s41593-022-01186-3.

