## Supplementary material for "From Belief to Brain: How Growth Mindset Optimizes Cortico-Striatal Dynamics for Cognitive Development"

**Examination of manipulation**

To verify the effectiveness of our WM-loading manipulation, we conducted repeated-measures analysis of variance (ANOVA) on behavioral performance with task loads as within-subject factors in all three years’ tests. At Year-1 visit, task loads showed robust main effects (*F*(1.94, 724.95) = 471.913, *p* < 0.001, *η^2^* = 0.308). Pairwise comparisons in the Year-2/3 revealed worse performance in higher loading condition (1- vs. 0-back: *t*(374) = -7.946; 2- vs. 1-back: *t*(374) = -23.075, both *p* < 0.001, Bonferroni-corrected; **Figure 1E**). In accordance with performance, participants responded slower as a function of WM loads (1- vs. 0-back: *t*(370) = 15.712; 2- vs. 1-back: *t*(367) = 11.415, both p < 0.001, Bonferroni-corrected; **Figure 1E**). Across all observations, workloads showed similar effect on WM performance **(Table S3 & S4)**.

**Longitudinal trajectory of WM**

We validated the reliability of N-back task and mindset measurement over time using intra-class correlations (ICCs) (**Table S5 & S6**). Based on the high effectiveness and reliability, we further investigated the longitudinal trajectory of WM performance using linear mixed model analysis. We compared a null model, a model with only gender effect, a linear age effect model and a quadratic age effect model (**Table S7 & S8**). The best fit models revealed a linear developmental trajectory of WM performance in all three WM loads (**Table S9 & S10**), with stronger effect on high task demands over three years. Thus, we took the linear effect of age into account in subsequent analyses.

**Comparison of meta-analytic and mindset-related maps**

We compared our whole-brain multiple regression results with the meta-analysis of WM from Neurosynth database (http://neurosynth.org). These two maps were highly overlapped, with conjunct clusters mainly distributed in dACC, right aIns, bilateral FEF and bilateral IPS (**Table S13A**). Considering the negative correlation found in parahippocampus and regions around PCC, we also compared the negative map of multiple regression analysis with meta-analysis of DMN. The result showed conjunct clusters in right parahippocampus, left calcarine and bilateral precuneus (**Table S13B**).

**Supplementary figures**

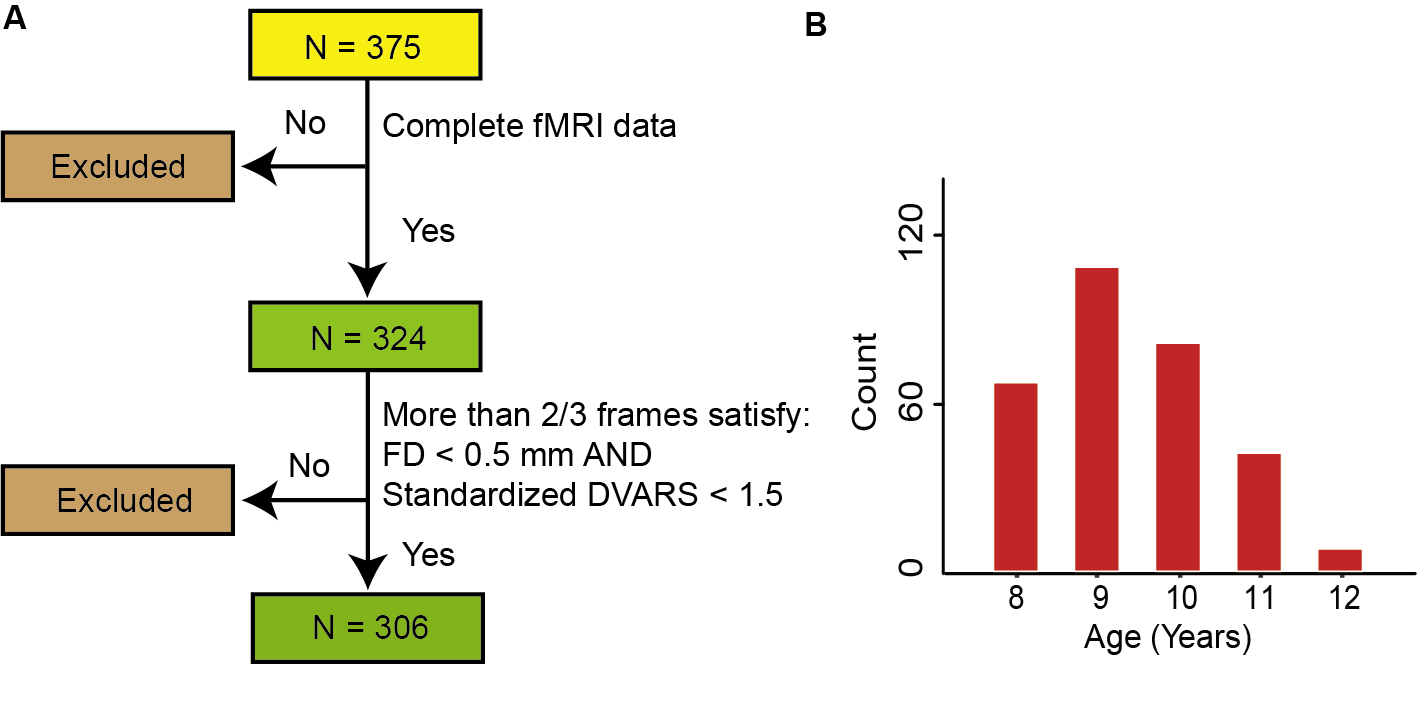

**Figure S1**. **Imaging data in Year-1 test. A.** Participant selection procedure for fMRI analysis. A total number of 375 children were included in the behavioral analysis. Children with incomplete data (n = 51) or excessive head motion during functional imaging (n = 18) were excluded from the fMRI analysis. The final sample size for fMRI analysis was 306. **B.** Age distribution of the final imaging sample (n = 306). Participants’ ages ranged from 8 to 12 years old in the first year.

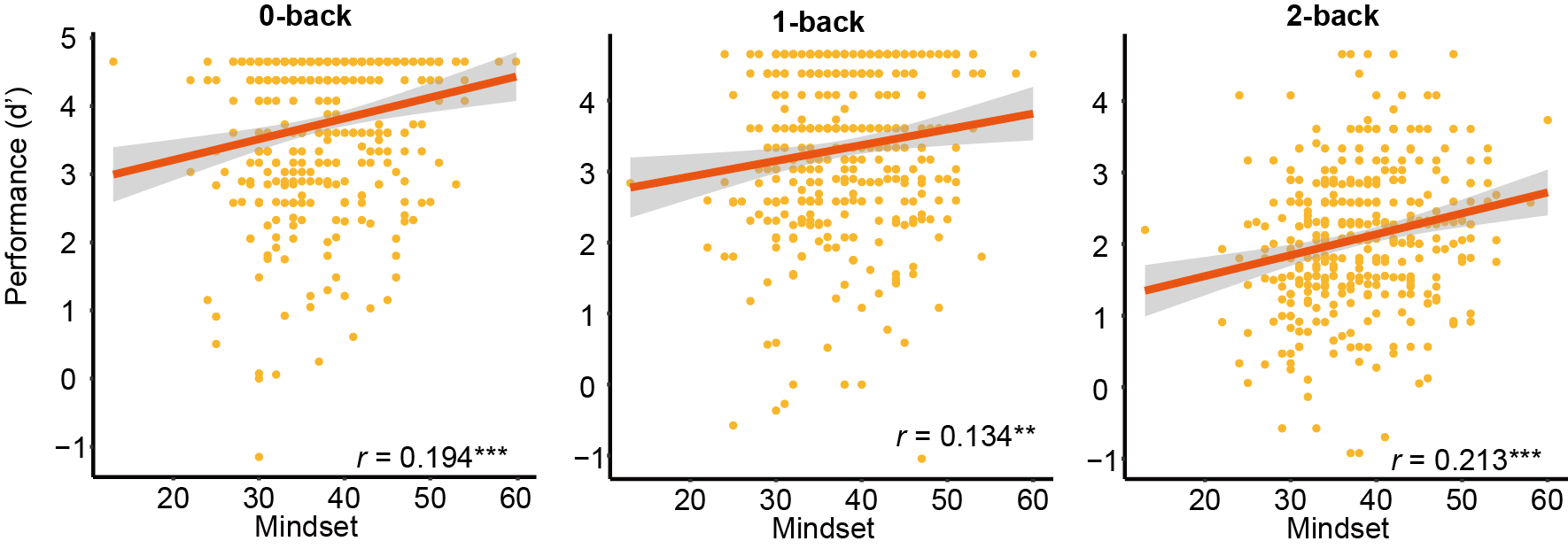

**Figure S2. Growth mindset positively predicted WM d’ in Year-1 test (n = 375).** Strongest effect of growth mindset was observed in 2-back condition. Age and gender was set as covariances. Each dot represents one child in one test. Shading represents 95% CI. ****p* < 0.001, ***p* < 0.01.

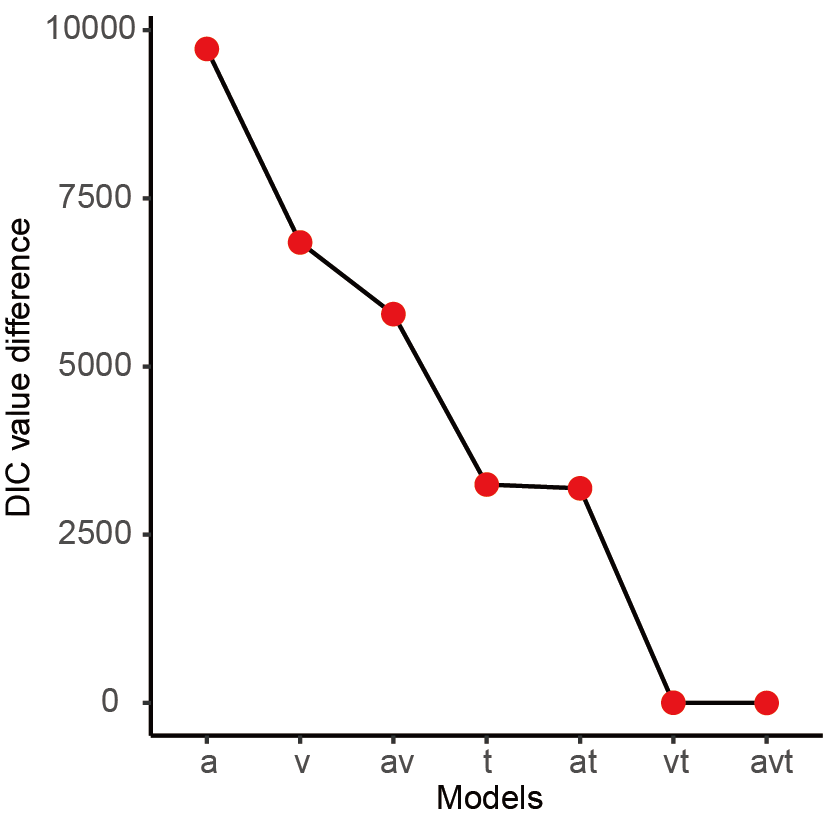

**Figure S3. Comparisons among 7 models.** Model comparisons were based on deviance information criterion (DIC), with lower DIC represents better model fitting. The y-axis represents DIC value differences between each alternative model and the best fit model (avt). The x-axis shows parameters of each model included for comparison. A total of 7 models with different free parameters were fitted for participants under three workloads across three years.

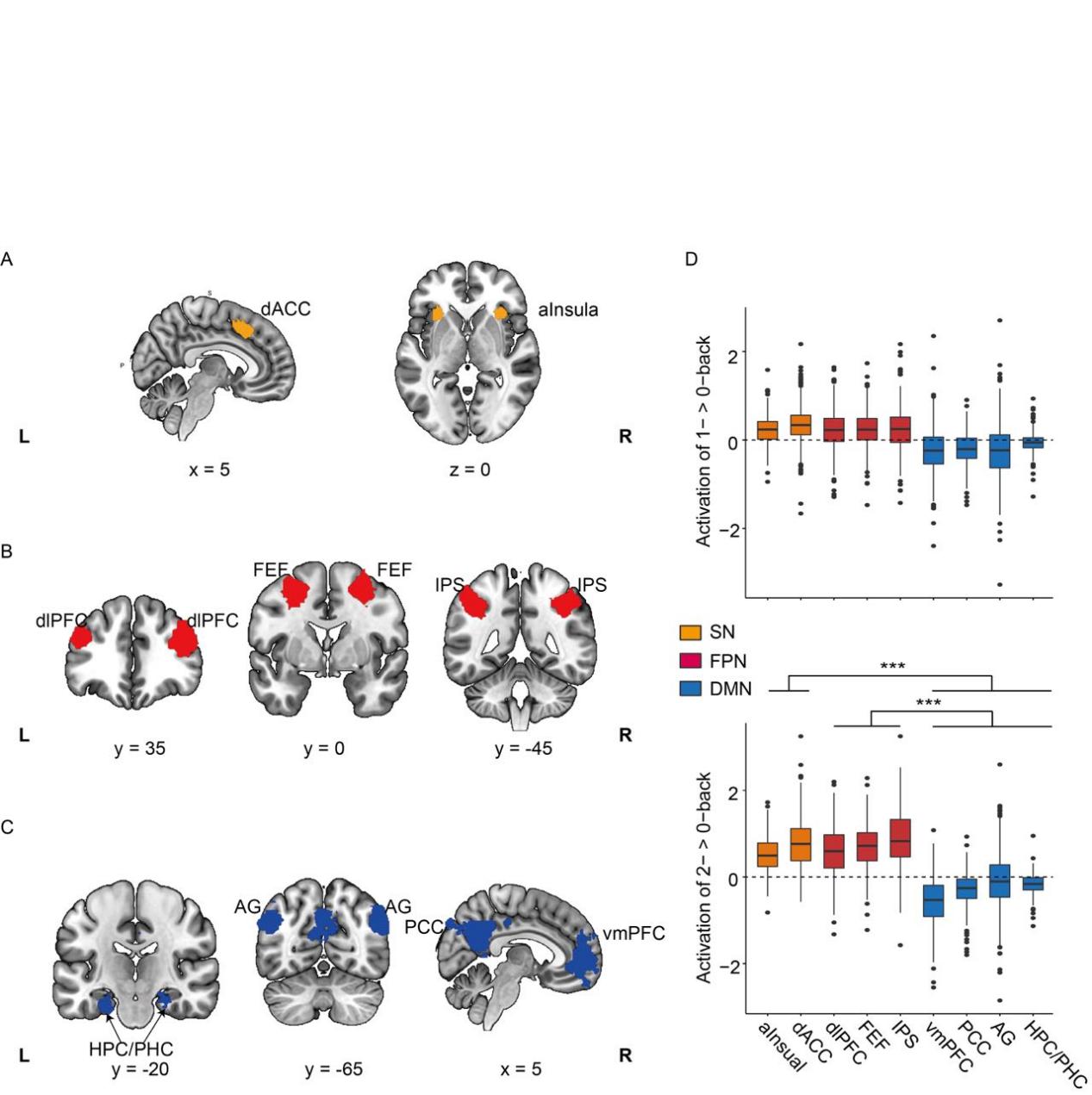

**Figure S4.** **ROI definition and task-related activation (n = 306).** **A.** ROIs defined for SN: dorsal anterior cingulate cortex (dACC); anterior Insula (aInsula). **B.** ROIs defined for FPN: dorsal lateral prefrontal cortex (dlPFC); frontal eye field (FEF); inferior parietal sulcus (IPS). **C.** ROIs defined for DMN: hippocampus/parahippocampus (HPC/PHC); angular gyrus (AG); posterior cingulate cortex (PCC); ventral medial prefrontal cortex (vmPFC). **D.** Activations of all ROIs in the contrasts of 1-back > 0-back and 2-back > 0-back. All ROIs from FPN and SN were activated, and all ROIs in DMN were dedactivated during the task. The thick black line in each box represents the median. The upper and lower edges of each box correspond to the 25th and 75th percentiles, respectively. The upper and lower error bars each represent the largest and smallest values within 1.5 times IQR (inter-quartile range, the distance between the 25th and 75th percentiles) above 75th percentile and below 25th percentile. ^***^*p* < 0.001.

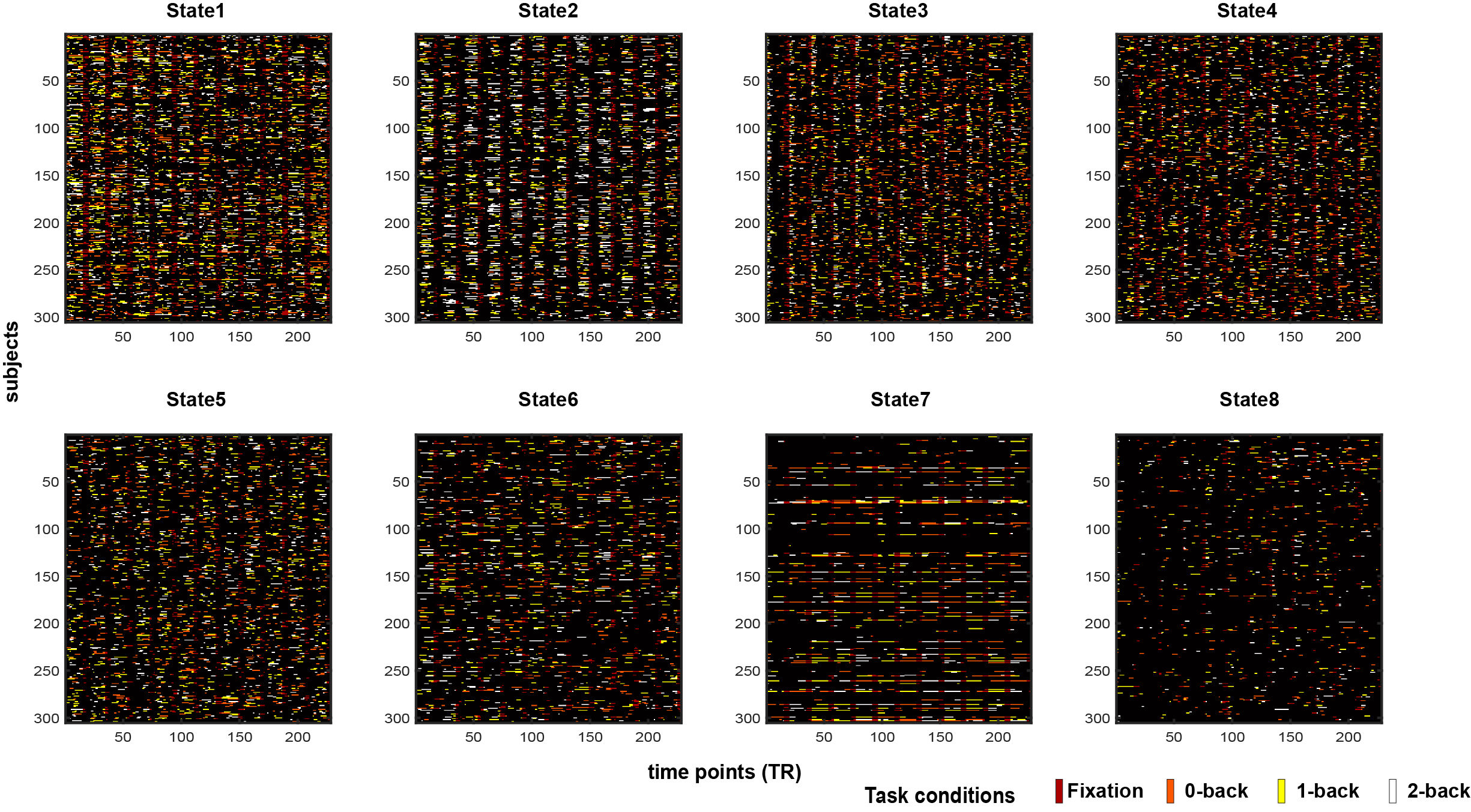

**Figure S5.** **The occurrences of states at each task conditions.** The presence of each state was binarized at each time point. If one state presented at a time point, the designed task condition would be labeled in the figure.

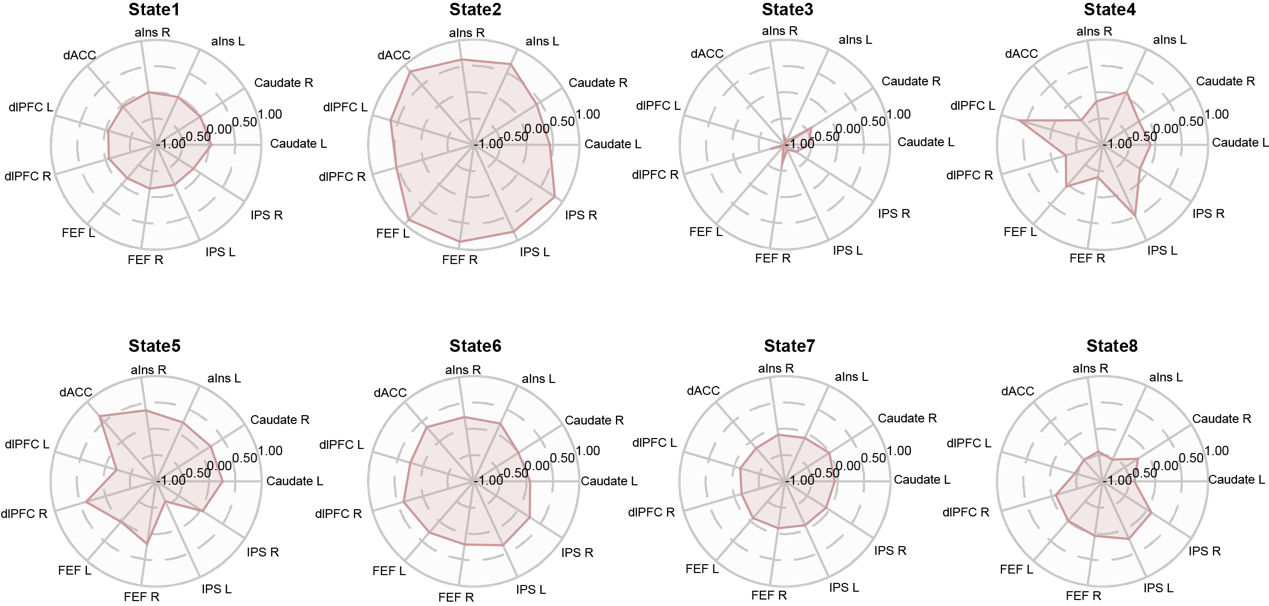

**Figure S6.** **Group-level spatial features of eight latent states.** Polar graphs of each brain state with the relative load of 11 cortico-striatal regions. The value of each area represents the relative activity magnitude referring to the mean activity.

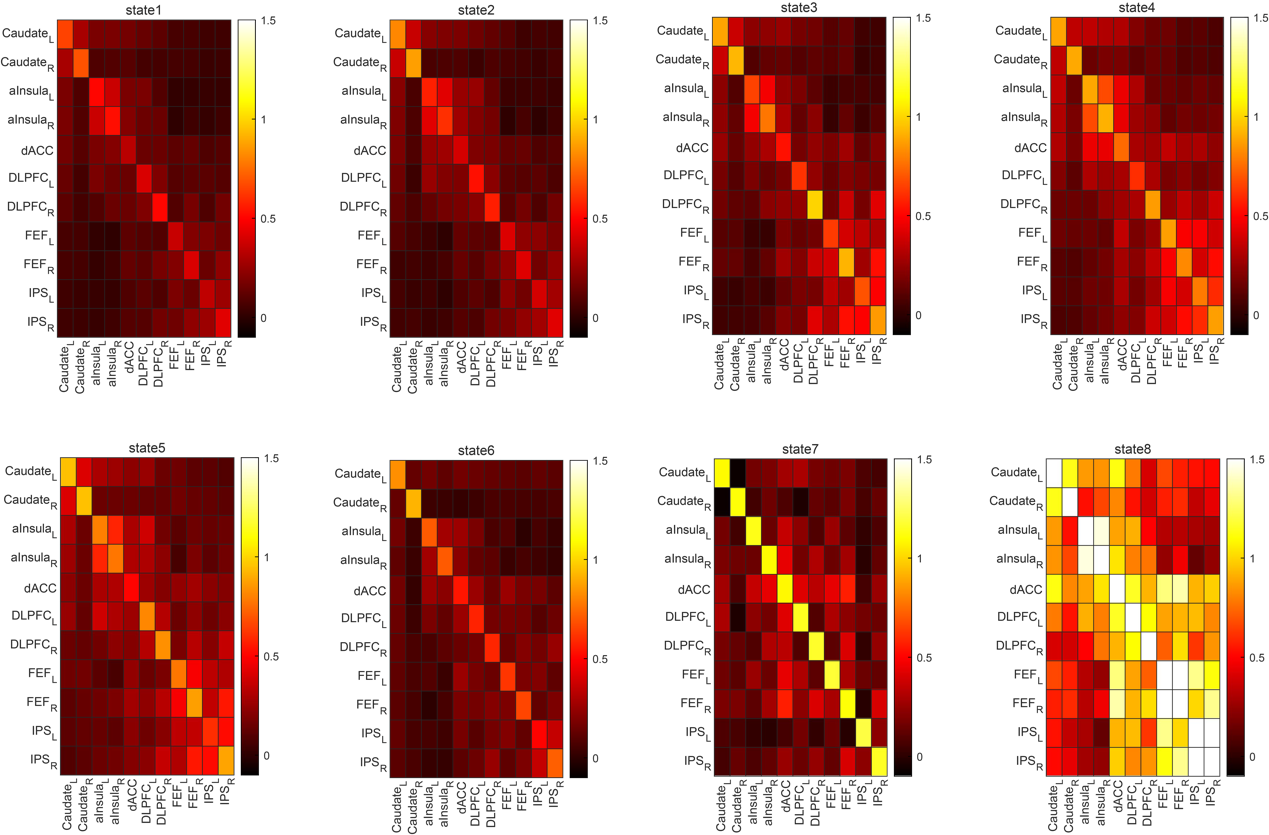

**Figure S7.** **Connectivity between ROIs for each latent state.** The covariance between ROIs in each state is presented in this plot.

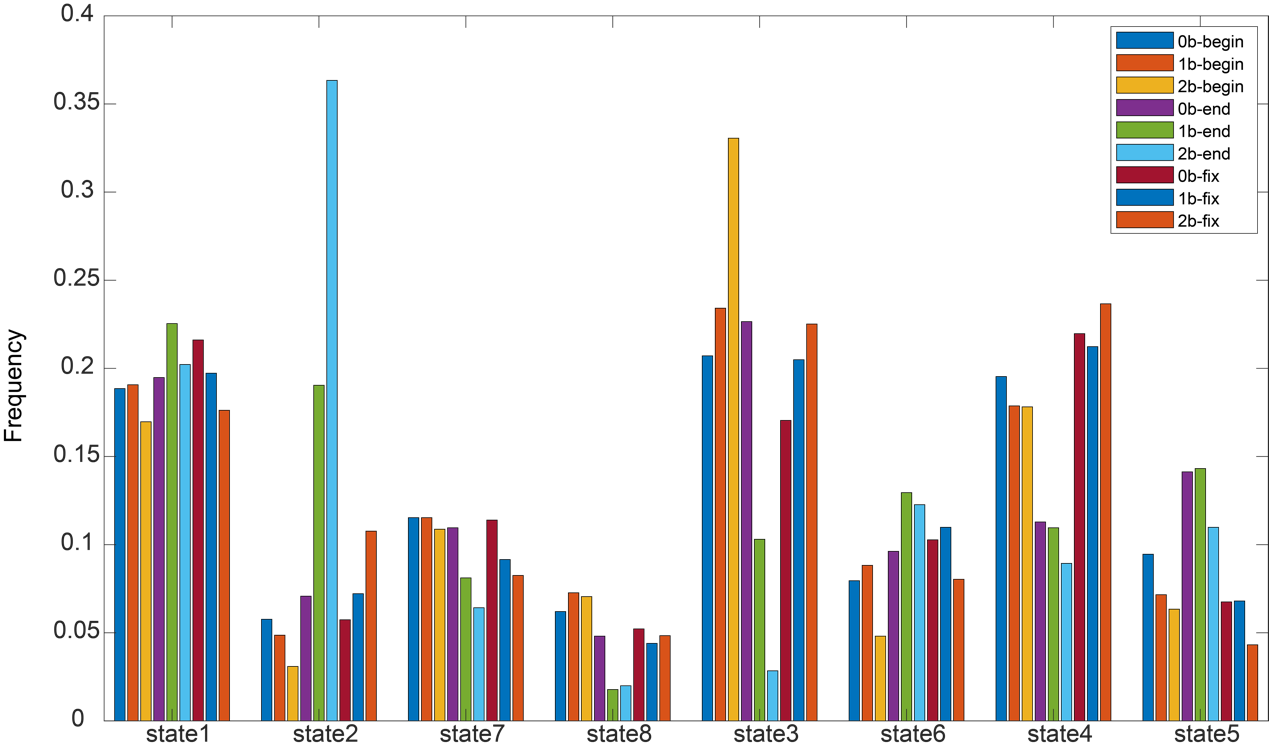

**Figure S8.** **Temporal distribution of each latent state during each block's beginning, end, and fixation.** The frequency of each state during the three frames (six seconds) at the beginning and end of each block, as well as during the fixation between blocks, was illustrated.

**Supplementary tables**

**Table S1**. Participant demographics of Year-1 and Year-2/3 tests

| **Analysis** | **Tests** | **n** | **Males/**  **Females** | **Age Range (years)** | **Average Age±SD (years)** |
| --- | --- | --- | --- | --- | --- |
| **Behavior** | **Year-1** | 375 | 207/168 | 8-12 | 9.833±1.039 |
|  | **Year-2** | 229 | 123/106 | 8-15 | 10.417±1.312 |
|  | **Year-3** | 144 | 69/75 | 8-13 | 10.944±1.223 |
| **Imaging** | **Year-1** | 306 | 163/143 | 8-12 | 9.869±1.024 |
|  | **Year-2/3** | 155 | 80/75 | 9-13 | 11.281±1.026 |

Notes: n, number of participants; SD, standard deviation.

**Table S2.** DIC values of model comparisons

| **Models** | **DIC** | **DIC_diff** |
| --- | --- | --- |
| a | 20627.41 | 9720.137 |
| v | 17751.34 | 6844.07 |
| av | 16685.81 | 5778.536 |
| t | 14153.24 | 3245.965 |
| at | 14096.98 | 3189.705 |
| vt | 10910.38 | 3.11166 |
| avt | 10907.27 | 0 |

Notes: a, decision threshold; v, drift rate; t, non-decision time

**Table S3**. Effect of workloads for WM performance

|  | **Year-1** | | **Year-2/3** | | | |
| --- | --- | --- | --- | --- | --- | --- |
|  | **n = 375** | | **Year-2 (n = 229)** | | **Year-3 (n = 144)** | |
|  | ***F*** | ***η2*** | ***F*** | ***η2*** | ***F*** | ***η2*** |
| d' | 471.91^***^ | 0.31 | 212.56^***^ | 0.26 | 105.75^***^ | 0.2 |
| RT | 325.86^***^ | 0.23 | 212.13^***^ | 0.2 | 111.54^***^ | 0.18 |

Notes: Main effect within each year were performed with repeated-measures analysis of variance (ANOVA). Some participants has no correct response in at least one of the conditions, which resulted in no RT records; so the total number of observation in Year1 for RT is 367. ****p* < 0.001.

**Table S4**. Effect of workloads for HDDM

|  | **Year-1** | | **Year-2/3** | | | |
| --- | --- | --- | --- | --- | --- | --- |
|  | **n = 375** | | **Year-2 (n = 229)** | | **Year-3 (n = 144)** | |
|  | ***F*** | ***η2*** | ***F*** | ***η2*** | ***F*** | ***η2*** |
| v | 917.00^***^ | 0.71 | 596.28^***^ | 0.72 | 351.258^***^ | 0.71 |
| a | 235093.37^***^ | 0.99 | 344837.50^***^ | 0.99 | 197521.13^***^ | 0.99 |
| t | 33.341^***^ | 0.08 | 16.12*** | 0.07 | 5.90** | 0.04 |

Notes: Main effect within each year were performed with repeated-measures analysis of variance (ANOVA). ****p* < 0.001, ** *p* < 0.01.

**Table S5**. Reliability of behavioral measurements over time.

| **Measure** | | | **ICC** |
| --- | --- | --- | --- |
| **Mindset (n = 80)** |  |  | 0.588*** |
| **N-back task (n = 78)** | d’ | 0-back | 0.610*** |
|  |  | 1-back | 0.604*** |
|  |  | 2-back | 0.756*** |
|  | RT | 0-back | 0.792*** |
|  |  | 1-back | 0.832*** |
|  |  | 2-back | 0.732*** |

Notes: Data were matched separately across three years for mindset and N-back task. Children with no correct response were excluded from the N-back matched data. ****p* < 0.001, ***p* < 0.01, Bonferroni-corrected.

**Table S6.** Reliability of HDDM parameters over time (n = 78)

| **Parameters** | | **ICC** |
| --- | --- | --- |
| a | 0-back | 0.408** |
|  | 1-back | 0.432** |
|  | 2-back | 0.561*** |
| v | 0-back | 0.511*** |
|  | 1-back | 0.596*** |
|  | 2-back | 0.643*** |
| t | 0-back | 0.718*** |
|  | 1-back | 0.779*** |
|  | 2-back | 0.591*** |

Notes: Data were matched separately across three years for mindset and N-back task. Children with no correct response were excluded from the N-back matched data. ****p* < 0.001, ***p* < 0.01, Bonferroni-corrected.

**Table S7.** Comparison of linear mixed models for d’ (N = 748)

| **Model** | **WM loads** | | | | | | **Mindset** | |
| --- | --- | --- | --- | --- | --- | --- | --- | --- |
|  | **0-back** | | **1-back** | | **2-back** | |  |  |
|  | **AIC** | **BIC** | **AIC** | **BIC** | **AIC** | **BIC** | **AIC** | **BIC** |
| **Null** | 2194.3 | 2208.1 | 2230.8 | 2244.7 | 2092.2 | 2106.1 | **4968.1** | **4982.0** |
| **Gender** | 2194.1 | 2212.5 | 2229.8 | 2248.2 | 2111.1 | 2110.7 | 4969.1 | 4987.6 |
| **Gender + age** | **2182.6** | **2205.7** | **2207.9** | **2231.0** | **1988.7** | **2011.8** | 4969.2 | 4992.3 |
| **Gender + age^2^** | 2184.4 | 2212.1 | 2209.9 | 2237.6 | 1990.7 | 2018.4 | 4966.6 | 4994.3 |

Notes: Best fitting models are indicated in bold. AIC: Akaike Information Criterion, BIC: Bayesian Information Criterion.

**Table S8. Comparison of linear mixed models for drift rate (N = 748)**

| **Model** | **WM loads** | | | | | |
| --- | --- | --- | --- | --- | --- | --- |
|  | **0-back** | | **1-back** | | **2-back** | |
|  | **AIC** | **BIC** | **AIC** | **BIC** | **AIC** | **BIC** |
| **Null** | 1589.1 | 1603.0 | 1445.5 | 1459.3 | 1473.2 | 1487.1 |
| **Gender** | 1588.0 | 1606.5 | 1445.3 | 1463.8 | 1472.5 | 1490.9 |
| **Gender + age** | **1527.6** | **1550.7** | **1392.6** | **1415.7** | **1384.0** | **1407.1** |
| **Gender + age^2^** | 1529.4 | 1557.1 | 1394.5 | 1422.2 | 1385.9 | 1413.7 |

Notes: Best fitting models are indicated in bold. AIC: Akaike Information Criterion, BIC: Bayesian Information Criterion.

**Table S9.** Linear mixed model of mindset and d’ over age and gender (N = 748)

| **Workloads** | **Model** | **Variance (*SD*)** | ***β*** | ***SE*** | ***t* value** | ***p*** |
| --- | --- | --- | --- | --- | --- | --- |
| 2-back | Random effects |  |  |  |  |  |
|  | Intercept | 0.336 (0.579) |  |  |  |  |
|  | Fixed effects |  |  |  |  |  |
|  | Intercept |  | 2.173 | 0.122 | 17.795 | <0.001 |
|  | Gender |  | 0.060 | 0.079 | 0.764 | 0.443 |
|  | Mindset |  | 0.122 | 0.034 | 3.580 | <0.001 |
|  | Age |  | 0.364 | 0.035 | 10.510 | <0.001 |
|  | Mindset *Age |  | -0.011 | 0.033 | -0.332 | 0.740 |
| 1-back | Random effects |  |  |  |  |  |
|  | Intercept | 0.251 (0.501) |  |  |  |  |
|  | Fixed effects |  |  |  |  |  |
|  | Intercept |  | 3.241 | 0.132 | 24.512 | <0.001 |
|  | Gender |  | 0.121 | 0.086 | 1.410 | 0.159 |
|  | Mindset |  | 0.119 | 0.039 | 3.027 | 0.003 |
|  | Age |  | 0.188 | 0.040 | 4.731 | <0.001 |
|  | Mindset *Age |  | -0.042 | 0.039 | -1.073 | 0.284 |
| 0-back | Random effects |  |  |  |  |  |
|  | Intercept | 0.329 (0.573) |  |  |  |  |
|  | Fixed effects |  |  |  |  |  |
|  | Intercept |  | 3.644 | 0.134 | 27.209 | <0.001 |
|  | Gender |  | 0.092 | 0.087 | 1.061 | 0.276 |
|  | Mindset |  | 0.120 | 0.039 | 3.101 | 0.002 |
|  | Age |  | 0.135 | 0.039 | 3.438 | <0.001 |
|  | Mindset *Age |  | -0.077 | 0.038 | -2.030 | 0.043 |

**Table S10.** Linear mixed model of mindset and drift rate over age and gender (N = 748)

| **Workloads** | **Model** | **Variance (*SD*)** | ***β*** | ***SE*** | ***t* value** | ***p*** |
| --- | --- | --- | --- | --- | --- | --- |
| 2-back | Random effects |  |  |  |  |  |
|  | Intercept | 0.126 (0.355) |  |  |  |  |
|  | Fixed effects |  |  |  |  |  |
|  | Intercept |  | 0.865 | 0.080 | 10.883 | <0.001 |
|  | Gender |  | 0.060 | 0.051 | 1.168 | 0.243 |
|  | Mindset |  | 0.087 | 0.023 | 3.853 | <0.001 |
|  | Age |  | 0.223 | 0.023 | 9.677 | <0.001 |
|  | Mindset *Age |  | -0.002 | 0.022 | -0.088 | 0.930 |
| 1-back | Random effects |  |  |  |  |  |
|  | Intercept | 0.093 (0.305) |  |  |  |  |
|  | Fixed effects |  |  |  |  |  |
|  | Intercept |  | 1.865 | 0.077 | 24.127 | <0.001 |
|  | Gender |  | 0.051 | 0.050 | 1.025 | 0.306 |
|  | Mindset |  | 0.091 | 0.023 | 3.987 | <0.001 |
|  | Age |  | 0.171 | 0.023 | 7.398 | <0.001 |
|  | Mindset *Age |  | -0.004 | 0.022 | -0.174 | 0.862 |
| 0-back | Random effects |  |  |  |  |  |
|  | Intercept | 0.104 (0.322) |  |  |  |  |
|  | Fixed effects |  |  |  |  |  |
|  | Intercept |  | 2.362 | 0.084 | 27.970 | <0.001 |
|  | Gender |  | 0.074 | 0.055 | 1.363 | 0.174 |
|  | Mindset |  | 0.059 | 0.025 | 2.359 | 0.020 |
|  | Age |  | 0.202 | 0.025 | 7.931 | <0.001 |
|  | Mindset *Age |  | -0.013 | 0.025 | -0.511 | 0.610 |

**Table S11**. Regions showed positive correlation with growth mindset (n = 306).

| **Region (AAL)** | | **Cluster size** | **t value** | **MNI coordinates** | | |
| --- | --- | --- | --- | --- | --- | --- |
|  |  |  |  | **X** | **Y** | **Z** |
| **2-back > 0-back** | **Parietal_Inf_L** | 904 | 5.2391 | -36 | -44 | 40 |
|  | **Supp_Motor_Area_L** | 362 | 4.9269 | -4 | 18 | 50 |
|  | **Frontal_Inf_Tri_R** | 99 | 4.8087 | 52 | 32 | 24 |
|  | **Parietal_Inf_R** | 667 | 4.6994 | 38 | -48 | 48 |
|  | **Precentral_L** | 553 | 4.5258 | -40 | 0 | 36 |
|  | **Frontal_Sup_2_R** | 149 | 4.4553 | 22 | 8 | 48 |
|  | **Precuneus_R** | 30 | 4.3207 | 8 | -74 | 54 |
|  | **Temoral_Pole_sup_R** | 12 | 4.2932 | 54 | 15 | -11 |
|  | **Putamen_R** | 24 | 4.2593 | 25 | 19 | 1 |
|  | **Precuneus_L** | 34 | 4.2185 | -6 | -70 | 60 |
|  | **Frontal_Mid_2_R** | 89 | 4.1872 | 48 | 24 | 38 |
|  | **Frontal_Mid_2_R** | 43 | 4.1347 | 38 | 52 | 6 |
|  | **Insula_R** | 73 | 4.085 | 38 | 18 | -2 |
|  | **Lingual_R** | 11 | 3.839 | 24 | -90 | -14 |
|  | **Precuneus_R** | 19 | 3.717 | 8 | -62 | 52 |
|  | **Precentral_L** | 17 | 3.6899 | -42 | 0 | 56 |
|  | **Caudate_L** | 23 | 3.6553 | -16 | -7 | 19 |
|  | **Calcarine_L** | 12 | 3.6128 | -2 | -86 | 0 |
|  | **Frontal_Mid_2_R** | 24 | 3.6113 | 36 | -2 | 62 |
|  | **Cingulum_Mid_R** | 24 | 3.603 | 8 | 36 | 30 |
|  | **Parietal_Sup_R** | 11 | 3.5637 | 20 | -70 | 58 |
|  | **Caudate_R** | 15 | 3.5139 | 16 | 4 | 20 |
|  | **Parietal_Sup_L** | 11 | 3.4508 | -18 | -66 | 56 |
|  | **Frontal_Inf_Oper_R** | 17 | 3.3216 | 44 | 16 | 30 |
|  | **Precentral_R** | 15 | 3.3103 | 32 | -6 | 46 |
| **2-back > 1-back** | **Supp_Motor_Area_L** | 120 | 4.7316 | -2 | 18 | 54 |
|  | **Parietal_Inf_L** | 353 | 4.5729 | -40 | -46 | 42 |
|  | **Occipital_Mid_L** | 32 | 4.4541 | -24 | -66 | 32 |
|  | **Frontal_Sup_2_R** | 22 | 4.2506 | 26 | 14 | 60 |
|  | **Temporal_Inf_L** | 10 | 4.101 | -44 | -52 | -12 |
|  | **Parietal_Inf_R** | 30 | 4.0815 | 34 | -56 | 50 |
|  | **Frontal_Sup_2_R** | 20 | 3.9752 | 22 | 8 | 48 |

Notes: One sample t-test was done for the coefficients of multiple regression. Clusters with peak t value passed the threshold of *p* < 0.05 (corrected for multiple comparison using FDR) were displayed. Cluster size: number of significant voxels of each cluster; t value: the peak t statistic of each continues cluster; MNI coordinates: corresponding MNI coordinates of the peak voxel. Only continues clusters with more than 10 voxels above threshold were presented. AAL: Automated Anatomical Labeling (AAL) template.

**Table S12**: Regions showed negative correlation with growth mindset (n = 306).

| **Region (AAL)** | | **Cluster size** | **t value** | **MNI coordinates** | | |
| --- | --- | --- | --- | --- | --- | --- |
|  |  |  |  | **X** | **Y** | **Z** |
| **2-back > 0-back** | **Calcarine_R** | 90 | -4.5585 | 18 | -50 | 10 |
|  | **ParaHippocampal_R** | 34 | -3.9588 | 28 | -36 | -14 |
|  | **ParaHippocampal_L** | 12 | -3.7805 | -32 | -38 | -10 |
|  | **Precuneus_L** | 27 | -3.5267 | -2 | -52 | 16 |
| **2-back > 1-back** | **Lingual_R** | 15 | -4.9462 | 14 | -54 | 6 |
|  | **Precuneus_L** | 24 | -4.6700 | -10 | -58 | 18 |
|  | **Calcarine_L** | 10 | -4.5242 | -6 | -58 | 4 |

Notes: One sample t-test was done for the coefficients of multiple regression. Clusters with peak t value passed the threshold of *p* < 0.001 in the contrast of 2-back > 0-back and *p* < 0.05 (FDR corrected) in the contrast of 2-back > 1-back were displayed. Cluster size: number of significant voxels of each cluster; t value: the peak t statistic of each continues cluster; MNI coordinates: corresponding MNI coordinates of the peak voxel. Only continues clusters with more than 10 voxels above threshold were presented. AAL: Automated Anatomical Labeling (AAL) template.

**Table S13:** Conjunction of meta-analysis and multiple regression in 2-back > 0-back contrast (n = 306)

| **Region (AAL)** | **Cluster size** | **MNI coordinates** | | |
| --- | --- | --- | --- | --- |
|  |  | **X** | **Y** | **Z** |
| A.    Positive results of multiple comparison & meta-analysis result of working memory | | | | |
| **Parietal_Inf_L** | 728 | -37 | -50 | 44 |
| **Parietal_Inf_R** | 529 | 38 | -49 | 45 |
| **Supp_Motor_Area_L** | 275 | -1 | 14 | 52 |
| **Precentral_L** | 249 | -43 | 7 | 33 |
| **Frontal_Sup_2_R** | 122 | 25 | 7 | 54 |
| **Precentral_L** | 110 | -28 | -4 | 55 |
| **Frontal_Inf_Tri_R** | 87 | 43 | 32 | 25 |
| **Precuneus_R** | 23 | 8 | -72 | 54 |
| **Insula_R** | 22 | 37 | 20 | -4 |
| **Precentral_R** | 18 | 47 | 9 | 35 |
| **Precuneus_R** | 15 | 9 | -62 | 51 |
| **Precuneus_L** | 12 | -8 | -71 | 58 |
| **Frontal_Sup_2_R** | 12 | 33 | -1 | 61 |
| **Frontal_Mid_2_R** | 11 | 36 | 50 | 8 |
| **Parietal_Sup_L** | 9 | -17 | -68 | 58 |
| **Precentral_R** | 8 | 32 | -6 | 48 |
| **Frontal_Mid_2_L** | 3 | -37 | 3 | 61 |
| **Caudate_L** | 1 | -16 | 0 | 18 |
| **Putamen_R** | 1 | 26 | 22 | -2 |
| **Frontal_Inf_Tri_R** | 1 | 44 | 22 | 30 |
| **Precuneus_L** | 1 | -4 | -66 | 56 |
| B.    Negative results of multiple comparison & meta-analysis result of DMN | | | | |
| **Precuneus_L** | 56 | -12 | -56 | 12 |
| **Precuneus_R** | 43 | 11 | -55 | 18 |
| **Cuneus_L** | 9 | -8 | -66 | 21 |
| **Calcarine_L** | 9 | -8 | -52 | 6 |
| **ParaHippocampal_R** | 7 | 28 | -34 | -14 |

Notes: Cluster size: number of significant voxels of each cluster; MNI coordinates: corresponding MNI coordinates of the cluster’s gravity center. AAL: Automated Anatomical Labeling (AAL) template.

**Table S14:** Conjunction of meta-analytic coactivation of striatum and multiple regression in 2-back > 0-back contrast (n = 306)

| **Region (AAL)** | **Cluster size** | **MNI coordinates** | | |
| --- | --- | --- | --- | --- |
|  |  | **X** | **Y** | **Z** |
| **Parietal_Inf_R** | 194 | 36 | -52 | 47 |
| **Parietal_Inf_L** | 119 | -45 | -41 | 45 |
| **Supp_Motor_Area_L** | 92 | 0 | 17 | 49 |
| **Frontal_Inf_Tri_R** | 64 | 44 | 33 | 25 |
| **Precentral_L** | 63 | -32 | -5 | 48 |
| **Precentral_L** | 54 | -40 | 0 | 34 |
| **Insula_R** | 46 | 38 | 19 | -3 |
| **Parietal_Inf_L** | 46 | -32 | -58 | 52 |
| **Caudate_L** | 23 | -16 | -7 | 19 |
| **Frontal_Sup_2_R** | 18 | 26 | 5 | 54 |
| **Caudate_R** | 15 | 17 | 1 | 19 |
| **Frontal_Inf_Oper_L** | 13 | -40 | 9 | 28 |
| **Supp_Motor_Area_L** | 13 | 0 | 5 | 63 |
| **Putamen_R** | 12 | 26 | 19 | 0 |
| **Frontal_Sup_2_R** | 12 | 34 | -2 | 61 |
| **Precentral_L** | 11 | -49 | 5 | 45 |
| **Precentral_R** | 10 | 33 | -5 | 51 |

Notes: Cluster size: number of significant voxels of each cluster; MNI coordinates: corresponding MNI coordinates of the cluster’s gravity center. AAL: Automated Anatomical Labeling (AAL) template.

**Table S15:** Partical correlation with mindset and WM performance of ROIs in the Year-1 test (n = 306)

| **ROIs** | | **Mindset** | | **Performance (d’)** | | **RT (ms)*** | |
| --- | --- | --- | --- | --- | --- | --- | --- |
|  |  | ***r*** | ***p*** | ***r*** | ***p*** | ***r*** | ***p*** |
| Striatum-CON | Striatum | 0.150 | 0.014 | 0.155 | 0.010 | 0.021 | 0.830 |
|  | aIns | 0.167 | 0.007 | 0.221 | <0.001 | 0.037 | 0.830 |
|  | SMA/dACC | 0.201 | 0.001 | 0.209 | <0.001 | 0.013 | 0.830 |
| FPN | dlPFC | 0.207 | 0.001 | 0.115 | 0.051 | -0.022 | 0.830 |
|  | FEF | 0.181 | 0.004 | 0.246 | <0.001 | -0.047 | 0.830 |
|  | IPS | 0.253 | <0.001 | 0.210 | <0.001 | 0.035 | 0.830 |
| DMN | vmPFC | -0.041 | 0.526 | -0.227 | <0.001 | -0.082 | 0.780 |
|  | PCC | -0.104 | 0.089 | -0.143 | 0.015 | -0.068 | 0.790 |
|  | AG | -0.033 | 0.563 | -0.094 | 0.101 | -0.091 | 0.780 |
|  | HPC/PHC | -0.117 | 0.060 | -0.217 | <0.001 | -0.018 | 0.830 |

Notes: Two-sided Pearson correlation partialed out age and gender effect, all p values were FDR corrected. Activation of each ROI were exacted from 2-back>0-back contrast.

*Three participants were excluded for RT analysis because of no correct response, results remains the same for d’ after removel of participants.

**Table S16:** Partical correlation with mindset and HDDM parameters of ROIs in the Year-1 test (n = 306)

| **ROIs** | | **Drift rate (v)** | | **Decision threshold (a)** | | **Non-decision time (t)** | |
| --- | --- | --- | --- | --- | --- | --- | --- |
|  |  | ***r*** | ***p*** | ***r*** | ***p*** | ***r*** | ***p*** |
| Striatum-CON | Striatum | 0.201 | <0.001 | 0.032 | 0.871 | 0.066 | 0.381 |
|  | aIns | 0.232 | <0.001 | 0.009 | 0.871 | 0.064 | 0.381 |
|  | SMA/dACC | 0.172 | 0.004 | -0.016 | 0.871 | 0.035 | 0.676 |
| FPN | dlPFC | 0.124 | 0.034 | -0.011 | 0.871 | -0.001 | 0.988 |
|  | FEF | 0.218 | <0.001 | -0.025 | 0.871 | 0.011 | 0.938 |
|  | IPS | 0.223 | <0.001 | 0.026 | 0.871 | 0.070 | 0.381 |
| DMN | vmPFC | -0.224 | <0.001 | -0.087 | 0.643 | -0.017 | 0.027 |
|  | PCC | -0.151 | 0.011 | -0.065 | 0.853 | -0.143 | 0.041 |
|  | AG | -0.096 | 0.096 | -0.096 | 0.643 | -0.149 | 0.041 |
|  | HPC/PHC | -0.157 | 0.009 | 0.022 | 0.871 | -0.097 | 0.225 |

Notes: Two-sided Pearson correlation partialed out age and gender effect, all p values were FDR corrected. Activation of each ROI were exacted from 2-back>0-back contrast.

**Table S17:** Mediation analysis of networks on the association between growth mindset and d’ in the Year-1 test (n = 306)

| **Striatum-CON** | | | |
| --- | --- | --- | --- |
|  | ***B*** | ***SE*** | **95% CI** |
| **a** | 0.211 | 0.054 | [0.106, 0.314] |
| **b** | 0.184 | 0.054 | [0.077, 0.289] |
| **c’** | 0.147 | 0.049 | [0.048, 0.240] |
| **ind** | 0.039 | 0.016 | [0.014, 0.077] |
| **FPN** | | | |
|  | ***B*** | ***SE*** | **95% CI** |
| **a** | 0.255 | 0.052 | [0.155, 0.357] |
| **b** | 0.171 | 0.056 | [0.062, 0.281] |
| **c’** | 0.142 | 0.050 | [0.043, 0.238] |
| **ind** | 0.043 | 0.017 | [0.015, 0.085] |

Notes: Path a, from X to M; path b, from M to Y; direct path c’, from X to Y; ind, indirect path a*b, indirect mediation effect of M; *SE*, standard error; 95% CI, 95% confidence interval. Age and gender were set as covariates. Activation of each ROI were exacted from 2-back>0-back contrast. All variables were Z transformed.

**Table S18:** Mediation analysis of networks on the association between growth mindset and drift rate (v) in the Year-1 test (n = 306)

| **Striatum-CON** | | | |
| --- | --- | --- | --- |
|  | ***B*** | ***SE*** | **95% CI** |
| **a** | 0.211 | 0.054 | [0.106, 0.314] |
| **b** | 0.167 | 0.056 | [0.059, 0.284] |
| **c’** | 0.127 | 0.052 | [0.024, 0.228] |
| **ind** | 0.035 | 0.016 | [0.011, 0.075] |
| **FPN** | | | |
|  | ***B*** | ***SE*** | **95% CI** |
| **a** | 0.255 | 0.052 | [0.155, 0.357] |
| **b** | 0.184 | 0.055 | [0.079, 0.294] |
| **c’** | 0.115 | 0.053 | [0.009, 0.217] |
| **ind** | 0.047 | 0.018 | [0.019, 0.089] |

Notes: Path a, from X to M; path b, from M to Y; direct path c’, from X to Y; ind, indirect path a*b, indirect mediation effect of M; *SE*, standard error; 95% CI, 95% confidence interval. Age and gender were set as covariates. Activation of each ROI were exacted from 2-back>0-back contrast. All variables were Z transformed.

**Table S19:** Partical correlation with mindset and Year-2/3 WM performance of ROIs (n = 155)

| **ROIs** | | **Mindset** | | **d’** | | **RT (ms)*** | |
| --- | --- | --- | --- | --- | --- | --- | --- |
|  |  | ***r*** | ***p*** | ***r*** | ***p*** | ***r*** | ***p*** |
| **Striatum-CON** | **Striatum** | 0.130 | 0.112 | 0.012 | 0.890 | -0.017 | 0.836 |
|  | **aIns** | 0.088 | 0.283 | 0.073 | 0.374 | -0.172 | 0.035 |
|  | **dACC** | 0.191 | 0.019 | 0.113 | 0.169 | -0.193 | 0.018 |
| **FPN** | **dlPFC** | 0.133 | 0.103 | 0.057 | 0.488 | -0.103 | 0.208 |
|  | **FEF** | 0.101 | 0.219 | 0.129 | 0.116 | -0.284 | <0.001 |
|  | **IPS** | 0.148 | 0.070 | 0.139 | 0.090 | -0.257 | 0.002 |
| **DMN** | **vmPFC** | -0.010 | 0.907 | -0.091 | 0.267 | 0.122 | 0.138 |
|  | **PCC** | -0.106 | 0.195 | -0.019 | 0.822 | -0.134 | 0.101 |
|  | **AG** | -0.053 | 0.520 | 0.034 | 0.680 | -0.124 | 0.131 |
|  | **HPC** | -0.063 | 0.444 | 0.012 | 0.887 | 0.049 | 0.549 |

Notes: Two-sided Pearson correlation partialed out age, gender and Year-1 performance. To present the differences between ROIs, all p values were not corrected for multiple comparison. Activation of each ROI were exacted from 2-back>0-back contrast.

*Two participants were excluded for RT analysis because of no correct response, results remains the same for d’ after removel of participants.

**Table S20:** Partical correlation with mindset and Year-2/3 HDDM parameters of ROIs (n = 155)

| **ROIs** | | **Drift rate (v)** | | **Decision threshold (a)** | | **Non-decision time (t)** | |
| --- | --- | --- | --- | --- | --- | --- | --- |
|  |  | ***r*** | ***p*** | ***r*** | ***p*** | ***r*** | ***p*** |
| Striatum-CON | Striatum | 0.030 | 0.711 | 0.130 | 0.109 | 0.007 | 0.929 |
|  | aIns | 0.099 | 0.225 | 0.064 | 0.433 | -0.124 | 0.128 |
|  | dACC | 0.199 | 0.014 | 0.063 | 0.441 | -0.066 | 0.416 |
| FPN | dlPFC | 0.157 | 0.053 | 0.076 | 0.352 | -0.010 | 0.904 |
|  | FEF | 0.172 | 0.034 | -0.073 | 0.374 | -0.159 | 0.050 |
|  | IPS | 0.130 | 0.110 | -0.018 | 0.823 | -0.154 | 0.059 |
| DMN | vmPFC | -0.104 | 0.203 | 0.111 | 0.174 | 0.054 | 0.513 |
|  | PCC | -0.062 | 0.449 | -0.004 | 0.961 | -0.183 | 0.024 |
|  | AG | -0.033 | 0.684 | -0.026 | 0.751 | -0.172 | 0.034 |
|  | HPC | -0.049 | 0.549 | 0.072 | 0.379 | -0.021 | 0.793 |

Notes: Two-sided Pearson correlation partialed out age and gender. To present the differences between ROIs, all p values were not corrected for multiple comparison. Activation of each ROI were exacted from 2-back>0-back contrast.

**Table S21:** Mediation analysis of striatum-CON on the association between growth mindset and 2-back d’ in the longitudinal subset (n =155)

| **Striatum-CON** | | | |
| --- | --- | --- | --- |
|  | ***B*** | ***SE*** | **95% CI** |
| **a** | 0.183 | 0.078 | [0.028, 0.355] |
| **b** | 0.188 | 0.073 | [0.045, 0.329] |
| **c’** | 0.079 | 0.079 | [-0.072, 0.239] |
| **ind** | 0.034 | 0.021 | [0.005, 0.090] |

Notes: Path a, from X to M; path b, from M to Y; direct path c’, from X to Y; ind, indirect path a*b, indirect mediation effect of M; *SE*, standard error; 95% CI, 95% confidence interval. Age and gender were set as covariates. Activation of each ROI was exacted from 2-back>0-back contrast. All variables were Z-transformed.

**Table S22:** Mediation analysis of striatum-CON on the association between growth mindset and drift rate (v) in the longitudinal subset (n = 155)

| **Striatum-CON** | | | |
| --- | --- | --- | --- |
|  | ***B*** | ***SE*** | **95% CI** |
| **a** | 0.183 | 0.078 | [0.028, 0.335] |
| **b** | 0.236 | 0.076 | [0.089, 0.388] |
| **c’** | 0.081 | 0.086 | [-0.086, 0.250] |
| **ind** | 0.043 | 0.024 | [0.008, 0.104] |

Notes: Path a, from X to M; path b, from M to Y; direct path c’, from X to Y; ind, indirect path a*b, indirect mediation effect of M; *SE*, standard error; 95% CI, 95% confidence interval. Age and gender were set as covariates. Activation of each ROI were exacted from 2-back>0-back contrast. All variables were Z transformed.

**Table S23**: Correlation between state frequency in whole task and children’s mindset and 2-back WM performance (n = 306)

|  |  | State1 | State2 | State3 | State4 | State5 | State6 | State7 | State8 |
| --- | --- | --- | --- | --- | --- | --- | --- | --- | --- |
| Mindset | *r* | 0.038 | **0.166** | **0.165** | 0.057 | 0.050 | -0.063 | **-0.142** | -0.052 |
|  | *p* | 0.505 | **0.016** | **0.016** | 0.436 | 0.436 | 0.436 | **0.035** | 0.436 |
| d' | *r* | 0.091 | **0.215** | **0.157** | 0.055 | -0.011 | -0.097 | **-0.137** | -0.141 |
|  | *p* | 0.153 | **0.001** | **0.024** | 0.391 | 0.843 | 0.149 | **0.034** | 0.034 |
| v | *r* | 0.079 | **0.185** | 0.114 | 0.063 | -0.014 | -0.079 | -0.116 | -0.128 |
|  | *p* | 0.228 | **0.010** | 0.093 | 0.309 | 0.803 | 0.228 | 0.093 | 0.093 |

Notes: Significant correlations (*p*<0.05) are marked in bold.

**Table S24**: Correlation between state flexibility and children’s mindset and 2-back WM performance (n = 306)

|  |  | State1 | State2 | State3 | State4 | State5 | State6 | State7 | State8 |
| --- | --- | --- | --- | --- | --- | --- | --- | --- | --- |
| Mindset | *r* | 0.112 | **0.191** | 0.082 | 0.005 | -0.015 | -0.070 | -0.125 | -0.088 |
|  | *p* | 0.135 | **0.007** | 0.244 | 0.932 | 0.910 | 0.299 | 0.119 | 0.244 |
| d' | *r* | **0.142** | **0.253** | 0.095 | -0.084 | -0.003 | -0.128 | **-0.142** | -0.097 |
|  | *p* | **0.036** | **<.001** | 0.133 | 0.166 | 0.955 | 0.053 | **0.036** | 0.133 |
| v | *r* | 0.132 | **0.218** | 0.094 | -0.071 | -0.053 | -0.090 | -0.116 | -0.075 |
|  | *p* | 0.085 | **0.001** | 0.191 | 0.247 | 0.360 | 0.191 | 0.116 | 0.247 |

Notes: Significant correlations (*p*<0.05) are marked in bold.

**Table S25**: Correlation between state stability and children’s mindset and 2-back WM performance (n = 306)

|  |  | State1 | State2 | State3 | State4 | State5 | State6 | State7 | State8 |
| --- | --- | --- | --- | --- | --- | --- | --- | --- | --- |
| Mindset | *r* | 0.019 | 0.100 | **0.161** | 0.006 | 0.111 | 0.055 | -0.006 | 0.138 |
|  | *p* | 0.918 | 0.167 | **0.040** | 0.918 | 0.146 | 0.539 | 0.918 | 0.066 |
| d' | *r* | 0.093 | **0.262** | **0.306** | 0.072 | 0.049 | 0.091 | **0.139** | **0.290** |
|  | *p* | 0.152 | **<.001** | **<.001** | 0.241 | 0.398 | 0.152 | **0.031** | **<.001** |
| v | *r* | 0.075 | **0.220** | **0.283** | 0.031 | 0.105 | 0.096 | 0.107 | **0.269** |
|  | *p* | 0.221 | **<.001** | **<.001** | 0.590 | 0.109 | 0.126 | 0.109 | **<.001** |

Notes: Significant correlations (*p*<0.05) are marked in bold.
